# ATF2 phosphorylation is a core transcriptional driver of neuron apoptosis

**DOI:** 10.1101/2023.09.27.559856

**Authors:** Jorge Gómez-Deza, Matthew Nebiyou, Lara H. El Touny, Mor R. Alkaslasi, Zhenyu Zuo, Josette J. Wlaschin, Francisco M. Nadal-Nicolas, Anastasia L. Slavutsky, Eliza Y. H. Lloyd, Peter M. Hayashi, Nathan Ashby, Mira Sohn, Ryan Dale, Wei Li, Ken Chih-Chien Cheng, Pedro P. Rocha, Claire E. Le Pichon

## Abstract

Apoptotic neuron death is a key feature of neurodegenerative disease. Considerable efforts have been made to target this pathway but the molecular mechanisms remain incompletely understood. Here, we conducted an unbiased whole genome CRISPR inhibition screen in human neurons to discover genes required for their death and identified known targets including the kinase MAP3K12 (DLK) and the transcription factor JUN. In addition, this screen revealed a potential role for the transcription factor ATF2. We demonstrate that ATF2 phosphorylation by MAP3 kinases is the core driver of the pro-apoptotic transcriptional response. Surprisingly, JUN phosphorylation is not required for apoptosis. However, the phosphorylation of ATF2 and upregulation of JUN expression are crucial. ATF2 therefore converts the kinase signal into a transcriptional response. Inhibiting ATF2 in cultured human neurons prevents cell death. Notably we show that ATF2 knockdown is neuroprotective in injury models *in vivo*. Thus, ATF2 provides a promising new target for a wide range of neurodegenerative disorders.

## Introduction

Neuron death is a key hallmark of neurodegenerative conditions such as Alzheimer’s disease (AD), amyotrophic lateral sclerosis (ALS) and glaucoma, as well as a common consequence of brain injury and chemotherapeutic treatment. As post-mitotic cells lacking stem cells, most neurons cannot be replaced. Neuron death can occur by different mechanisms including apoptosis, autophagy, ferroptosis, necrosis, and others^1^. Among these, apoptosis has been the most extensively studied form of cell death and is mediated by the caspase protein family. However, the precise mechanisms gating apoptosis and by which apoptosis occurs in neurons remain unclear. A better understanding of the molecular steps leading to neuron death would have significant value for patients suffering from diverse neurodegenerative conditions, from traumatic brain injury or chemotherapy-induced peripheral neuropathy to AD or ALS.

Dual leucine zipper kinase (DLK or MAP3K12) is an evolutionarily conserved, neuronally enriched MAP triple kinase and a key regulator of axon degeneration and neuron death. Leucine zipper kinase (LZK or MAP3K13) is a highly homologous but less well-studied MAP triple kinase, sharing catalytic and leucine zipper domains with DLK^2^. Much work has focused on DLK, whereas the relative importance of LZK in neuron degeneration and regeneration has only recently been examined^3–7^. Moreover, how DLK and LZK cooperate to regulate neuronal death in human neurons remains unknown. The DLK pathway has been recognized as a therapeutic target for many neurodegenerative conditions, prompting the development of DLK inhibitors^8–11^. However, a clinical trial of DLK inhibitors in ALS patients was halted due to safety concerns^12^, highlighting a need to better understand the DLK/LZK pathway and identify alternate candidate targets.

To discover critical factors contributing to neuronal apoptosis, we performed whole-genome CRISPR interference (CRISPRi) screen on human neurons treated with vincristine. This chemotherapeutic agent drives DLK-dependent neurotoxicity through destabilization of microtubules. Using this system, we identified DLK, JUN, and activating transcription factor 2 (ATF2), among others, as essential regulators of apoptosis in human neurons. While DLK and JUN are known players in this pathway, much less is known about ATF2.

DLK regulates the neuronal transcriptional response to stress and injury via the phosphorylation of MAP2K4/7 and Jun N-terminal Kinase (JNK), which lead to the phosphorylation and activation of several transcription factors including cJun (the protein encoded by the *JUN* gene)^13^. cJun is regarded as a critical regulator of neurodegeneration by activating the transcription of pro-apoptotic factors. Whether cJun phosphorylation is necessary to drive apoptosis in neurons is still unclear^14–18^ and a question we address in this study. cJun is a member of the activator protein-1 (AP-1) family of transcription factors, characterized by their basic leucine zipper (bZip) domain. Additional AP-1 proteins include cFOS, ATF4, ATF3 and ATF2, which homo- and heterodimerize via their bZip domains to regulate transcription^19^. Activating transcription factors (ATF) are a subfamily of the AP-1 transcription factors that share preferential binding to cAMP responsive elements (CRE). They include 7 members, ATF1- 7^20^. Although ATF3 and ATF4 are well characterized contributors to neuronal injury responses^21–24^, the role of ATF2 in neurons has been largely overlooked.

Studies in cell lines have revealed ATF2 can be phosphorylated by several proteins, driving its nuclear localization. Once in the nucleus, it exerts transcriptional activity by homo- and heterodimerization with cJun^25^. ATF2 is highly expressed in the brain and its depletion causes cranial motor neuron degeneration^26^, highlighting its essential function in neurodevelopment.

Here, we show that DLK and LZK cooperate to induce neuron death by regulating the pro-apoptotic transcriptional response in human neurons. Each of these kinases is necessary for neuron death. The phosphorylation of cJun, a reliable marker of DLK pathway activation, has been thought of as a critical step for neuron death downstream of DLK. However, we find that cJun phosphorylation is dispensable for vincristine-induced neurodegeneration, and rather that its transcriptional upregulation is key. Our experiments identify ATF2 phosphorylation as an essential driver of neurodegeneration upstream of JUN that links DLK/LZK signaling to JUN transcription. We demonstrate that this core function of phospho-ATF2 in driving neuron death is a highly conserved mechanism, from human neurons exposed to cytoskeletal disruption *in vitro* to two different *in vivo* mouse models of neuron injury. This study highlights the potential of targeting ATF2 phosphorylation to prevent neuron death across a variety of neurodegenerative conditions.

## Results

### Vincristine-induced neurodegeneration is transcription-dependent

To investigate the molecular mechanisms that lead to cell death in human neurons, we used a transcription factor-mediated platform to differentiate cortical- (NGN2-driven^27^ or i^3^ iPSCs) and sensory-like (NGN2/BRN3A-driven^28^) neurons from iPSCs (Fig. 1A). We replicated experiments in each of these two human neuron subtypes to increase confidence in our results. Importantly, both types of neurons are transcriptionally similar to human neurons from postmortem tissue^28,29^.

**Figure 1:**
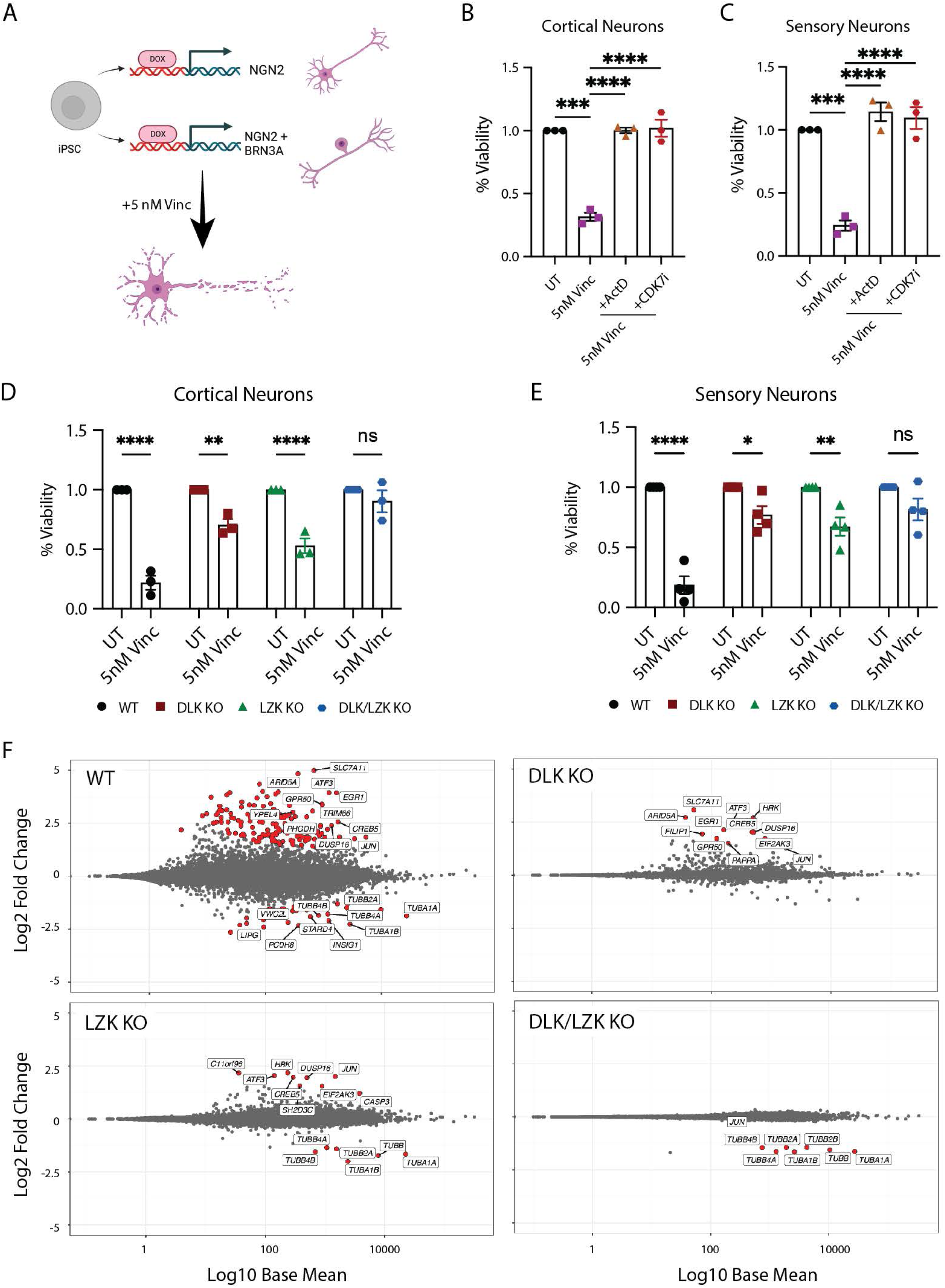
Vincristine-induced cell death is transcriptionally controlled by DLK and LZK A. Schematic representation of cortical and sensory i^3^Neuron differentiation generated using Biorender. B. Quantification of the relative viability of cortical i^3^Neurons after 48 hours of treatment with 5 nM vincristine, 5 nM vincristine + 30 nM ActD or 5 nM vincristine + 100 nM CDK7i. N=3 independent differentiations. Statistics for B and C: one-way ANOVA, Bonferroni correction, ns = not significant, p<0.01 **, p<0.005 ***, p<0.001 ****. C. Quantification of the relative viability of sensory i^3^Neurons after 48 hours of treatment with 5 nM vincristine, 5 nM vincristine + 30 nM ActD or 5 nM vincristine + 100 nM CDK7i. N=3 independent differentiations. D. Quantification of the relative viability of WT, DLK KO, LZK KO and double DLK/LZK KO cortical i^3^Neurons after 48 hours of 5 nM vincristine. N=4 independent differentiations. Statistics for D and E: two-way ANOVA, Bonferroni correction, ns = not significant, p<0.05 *, p<0.01 **, p<0.001 ****. E. Quantification of the relative viability of WT, DLK KO, LZK KO and double DLK/LZK KO sensory i^3^Neurons after 48 hours of 5 nM vincristine. N=3 independent differentiations. F. MA plots showing the transcriptional changes in WT, DLK KO, LZK KO and double DLK/LZK KO cortical i^3^Neurons after 16 hours of 5 nM vincristine. Genes that are significantly differentially expressed are labelled in red.

We first asked whether vincristine-induced neuronal death depends on transcription. Cell viability was assessed using an MTT assay in neurons treated for 48 hours with 5 nM vincristine with or without a transcriptional inhibitor, Actinomycin D (ActD) or cyclin dependent kinase 7 (CDK7) inhibitor, THZ1 2HCL (CDK7i). Vincristine treatment resulted in a significant reduction in neuron viability, which was rescued by inhibiting transcription using either ActD or CDK7i (Fig. 1B and C). Vincristine-induced neurodegeneration is therefore transcriptionally driven.

### DLK and LZK cooperate to regulate cell death in human neurons

DLK and LZK are highly similar neuronally-enriched stress-sensing proteins. They function by relaying axonal stress and initiating a transcriptional response that regulates the neuronal fate to injury. Despite high homology in their functional domains, relatively little is known about the role of LZK and how it may regulate transcription after neuronal stress and its role in regulating transcription. This high degree of similarity in their kinase domains of DLK and LZK make it impossible to specifically target either of them using small molecule inhibitors. Therefore, to assess their individual roles in the stress response to vincristine we generated DLK KO (knockout), LZK KO, and DLK/LZK double KO iPSCs and differentiated them into cortical and sensory neurons. Validation of DLK and LZK knockout was performed by PCR of the excised allele and by western blotting (Supp. Fig. 1A-F).

We verified whether DLK and LZK regulate cell death in human neurons in response to vincristine exposure by assessing the viability of wildtype (WT), DLK KO, LZK KO and DLK/LZK double KO cortical and sensory neurons treated with 5nM vincristine for 48 hours. WT, DLK KO and LZK KO, but not DLK/LZK KO cortical neurons showed a significant decrease in viability in response to vincristine exposure (Fig. 1D). We observed similar results in sensory neurons, where the deletion of both DLK and LZK fully protected neurons from degeneration, while individual KO of DLK or LZK was only partially protective (Fig. 1E).

Additionally, we measured the levels of phosphorylated H2AX (γH2AX), a marker of DNA damage that accumulates in the nucleus during cell death^30^. Treatment with vincristine for 24 hours resulted in a significant increase in the levels of γH2AX in WT neurons. Both the DLK and LZK KO lines significantly prevented γH2AX accumulation after vincristine treatment, while the double DLK/LZK KO completely abolished this response (Supp. Fig. 1G and H). We thus conclude that DLK and LZK cooperate to regulate vincristine-induced neuron death.

### DLK and LZK regulate the transcriptional response to vincristine

Since we had demonstrated that vincristine-induced neurodegeneration is transcription-dependent, and that DLK and LZK both regulate neuronal death, we set out to identify the DLK-, LZK- and DLK/LZK-dependent transcriptional changes induced by vincristine. We therefore treated WT, DLK, LZK and DLK/LZK KO neurons with vincristine for 16 hours and performed RNA sequencing (RNAseq). We chose this time point because inhibiting transcription up to 8 hours after vincristine addition prevented cell death, suggesting that essential transcriptional changes would be detected at a later timepoint (Supp Fig. 2A and B). We identified 154 differentially expressed genes in WT neurons after vincristine treatment (Fig. 1F and Supp. Fig 2C). In contrast, only 9 and 11 genes were differentially expressed in DLK and LZK KO neurons respectively, and almost no transcriptional response to vincristine was observed in double DLK/LZK KO neurons (Fig. 1F, Supp. Fig. 2C and Supp. Table 1). Gene ontology analysis of significantly upregulated genes in WT neurons treated with vincristine revealed enrichment of genes involved in the regulation of apoptosis and regulation of endoplasmic reticulum stress in WT neurons in response to vincristine (Supp. Fig. 2D). Little, if any, upregulation of these pathways was observed in all 3 KO lines after vincristine treatment (Supp. Fig 2E and F). Interestingly, DLK and LZK-dependent genes overlapped greatly (Supp. Fig. 2G), indicating that these proteins play redundant roles in regulating transcription in human neurons.

**Figure 2:**
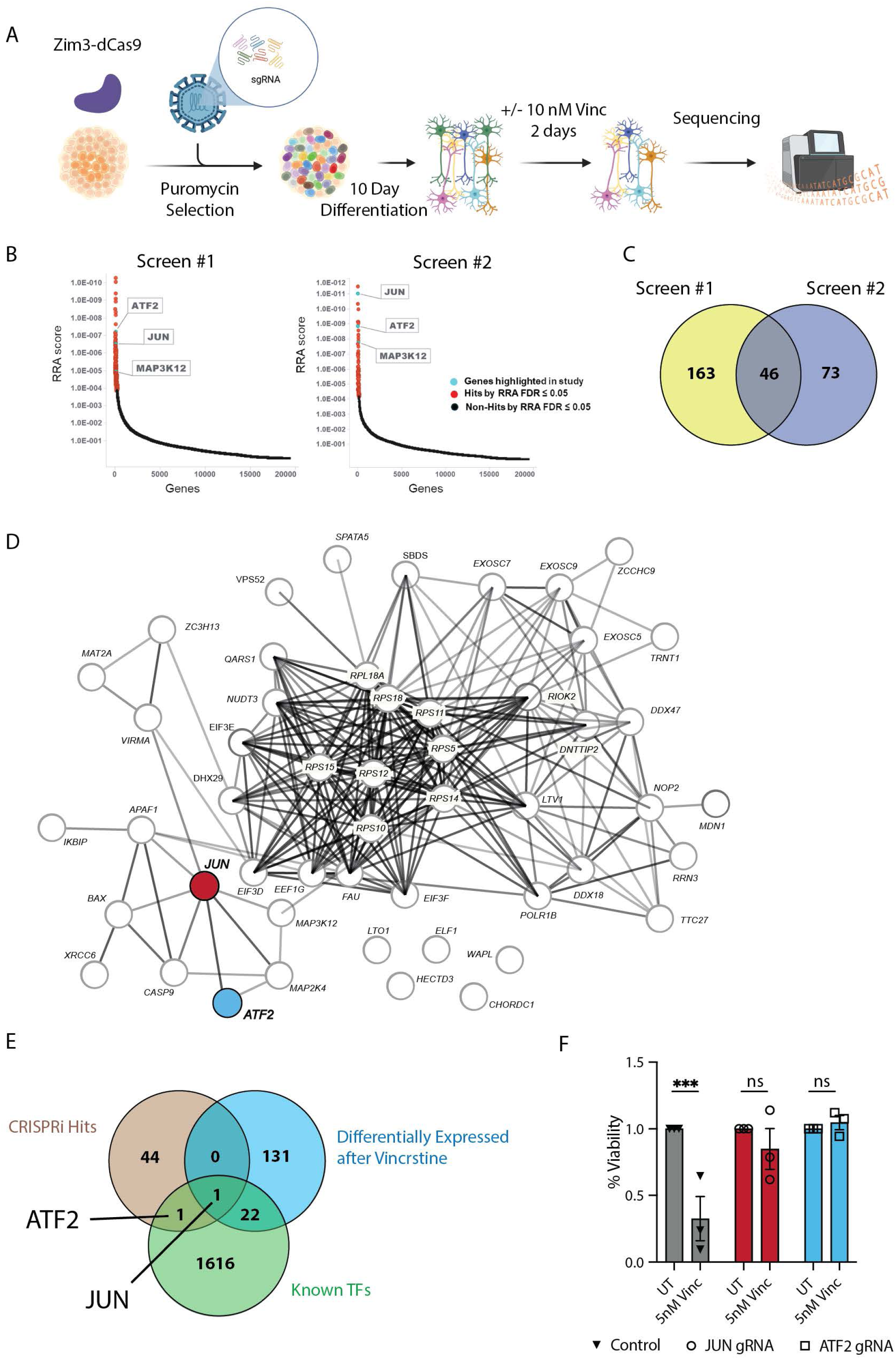
Genome-wide CRISPRi screen identifies cJun and ATF2 as regulators of cell death A. Schematic representation of the whole genome CRISPRi survival screen to identify modifiers of neuronal apoptosis in human neurons. Schematic made using Biorender. B. Top genes identified to promote neuron survival in screens 1 and 2. RRA scores in the positive selection direction in vincristine-treated vs untreated samples (lower depicted score indicates a stronger confidence in increasing survival when repressed in vincristine treated samples). C. Venn diagram showing genes common to both whole genome CRISPRi survival screens that significantly prevent vincristine-induced neuronal apoptosis. False discover rate (FDR) ≤0.05. D. Protein-protein interaction network for hits identified to suppress vincristine-induced neuronal apoptosis in two whole genome CRISPRi survival screens. *ATF2* and *JUN* are highlighted in red and blue respectively. E. Venn diagram showing the intersection of genes upregulated in WT i^3^Neurons after vincristine treatment by RNA seq (blue), genes identified to protect neurons from vincristine-induced neurodegeneration in i^3^Neurons (beige) and known transcription factors (green). F. Quantification of the relative viability of control, JUN and ATF2 knockdown cortical i^3^Neurons after 48 hours of 5 nM vincristine. Results are represented as mean ± SEM. N=3 independent differentiations. Two-way ANOVA, Bonferroni correction, ns = not significant, p<0.005 ***.

### Whole genome CRISPRi screen identifies JUN and ATF2 as upstream modifiers of neuronal apoptosis

To identify additional factors driving neurodegeneration in human neurons, we performed a whole genome CRISPR-interference (CRISPRi) screen. I^3^ iPSCs containing Zim3- dCas9 gene silencing machinery were transduced with a pooled library of dual gRNAs targeting all genes in the genome, as well as 2500 non-targeting controls. iPSCs transduced with gRNAs were selected using puromycin and differentiated into mature cortical neurons for 10 days. Neurons were then treated with 10 nM vincristine for 48 hours, or DMSO as a control. Next generation sequencing was performed on the surviving neurons in both conditions to identify genes that promoted neuronal survival after vincristine treatment (Fig. 2A).

The CRISPRi screen was performed twice, using slightly different cell collection methods and increasing reliability of our results (Fig. 2B, see methods for details). Genes with a false discovery rate (FDR) of ≤0.05 were considered positive hits. Screens 1 and 2 identified 209 and 119 genes respectively, and the intersection of both screens revealed 46 common genes that protect neurons against vincristine-induced neurotoxicity (Fig. 2C and Supp. Table 2). STRING-DB was used to create a protein-protein interaction network of genes driving neurotoxicity (Fig. 2D). Further gene ontology analysis highlighted the involvement of these proteins in ribosomal biogenesis, cytoplasmic translation as well as regulation of neuron apoptosis (Supp. Fig. 3A). Interestingly, we found little overlap between the genes transcriptionally upregulated by vincristine and the genes that were protective in our CRISPRi screen (Fig. 2E) with the sole exception of *JUN* (which encodes cJun) (Fig. 2D, red and 2E). Our results also identified a relatively understudied *JUN* interactor, the activating transcription factor-2 (*ATF2*) (Fig. 2D, blue and 2E).

We validated the CRISPRi screen hits *JUN* and *ATF2* by transducing Zim3-dCas9 iPSCs with gRNAs targeting each gene or a control gRNA and differentiated them into mature neurons. We confirmed the knockdown (KD) of these proteins by Western blotting (Supp. Fig. 3B-D). Interestingly, we found that ATF2 KD resulted in a significant reduction of total cJun (Supp. Fig. 3B-D). Silencing either JUN and ATF2 protected neurons from vincristine-induced neurotoxicity, validating these proteins as true positive hits in the screen (Fig. 2F).

### DLK and LZK regulate total and phospho-cJun levels in human cortical neurons via JNK and transcription

Phosphorylation of the transcription factor cJun (p-cJun) is widely regarded as a key regulatory step in DLK-dependent transcriptional responses^13^. We performed immunofluorescence on vincristine-treated WT, DLK KO, LZK KO and DLK/LZK KO neurons to measure the relative contribution of DLK and LZK to p-cJun levels. Interestingly, unlike in mouse neurons where cJun phosphorylation is almost entirely dependent on DLK alone^31–33^, deletion of either DLK or LZK alone was not sufficient to prevent phosphorylation of cJun in human neurons. Only the combined deletion of both DLK and LZK completely blocked cJun phosphorylation in human neurons treated with vincristine (Supp. Fig. 4A and B). To better understand the roles of these kinases in cJun regulation we also measured the levels of total cJun in the nucleus. Similarly, the increase in total cJun levels after vincristine was only partially decreased in single DLK or LZK KO neurons and prevented only in double KO neurons (Supp. Fig. 4C). Together these results mirror our RNAseq data, where JUN transcription is elevated in WT, DLK KO and LZK KO but not in double DLK/LZK KO neurons (Supp. Fig. 4D and Supp. Table 1).

To better understand how cJun and its phosphorylation are regulated, we treated WT neurons with vincristine and GNE-3511, a DLK/LZK inhibitor (DLKi), a Jun N-terminal Kinase (JNK) inhibitor (JNKi), or one of two transcription inhibitors (ActD and CDK7i). Treatment with any of these four inhibitors completely suppressed p-cJun increase after vincristine (Fig. 3A and B), indicating that the phosphorylation of cJun is regulated by the DLK/LZK-JNK pathway as well as by transcription. Additionally, total cJun levels were significantly decreased in neurons treated with any of the 4 inhibitors (Fig. 3C). This suggests that the observed increase in p-cJun levels may be due to a corresponding increase in total cJun levels. To test this, we performed double immunolabelling using an antibody recognizing p-cJun (S63) and a fluorescently conjugated antibody recognizing total cJun. Using this method, we correlated the levels of both protein species in the nucleus of individual cells before and after vincristine treatment. As expected, the levels of cJun and p-cJun (S63) highly correlated both before (black squares, Pearson correlation r=0.64 p<0.0001) and after (red triangles, Pearson correlation r=0.89 p<0.0001) vincristine treatment, indicating that phospho- and total cJun both increase under stress (Fig. 3D and E).

**Figure 3:**
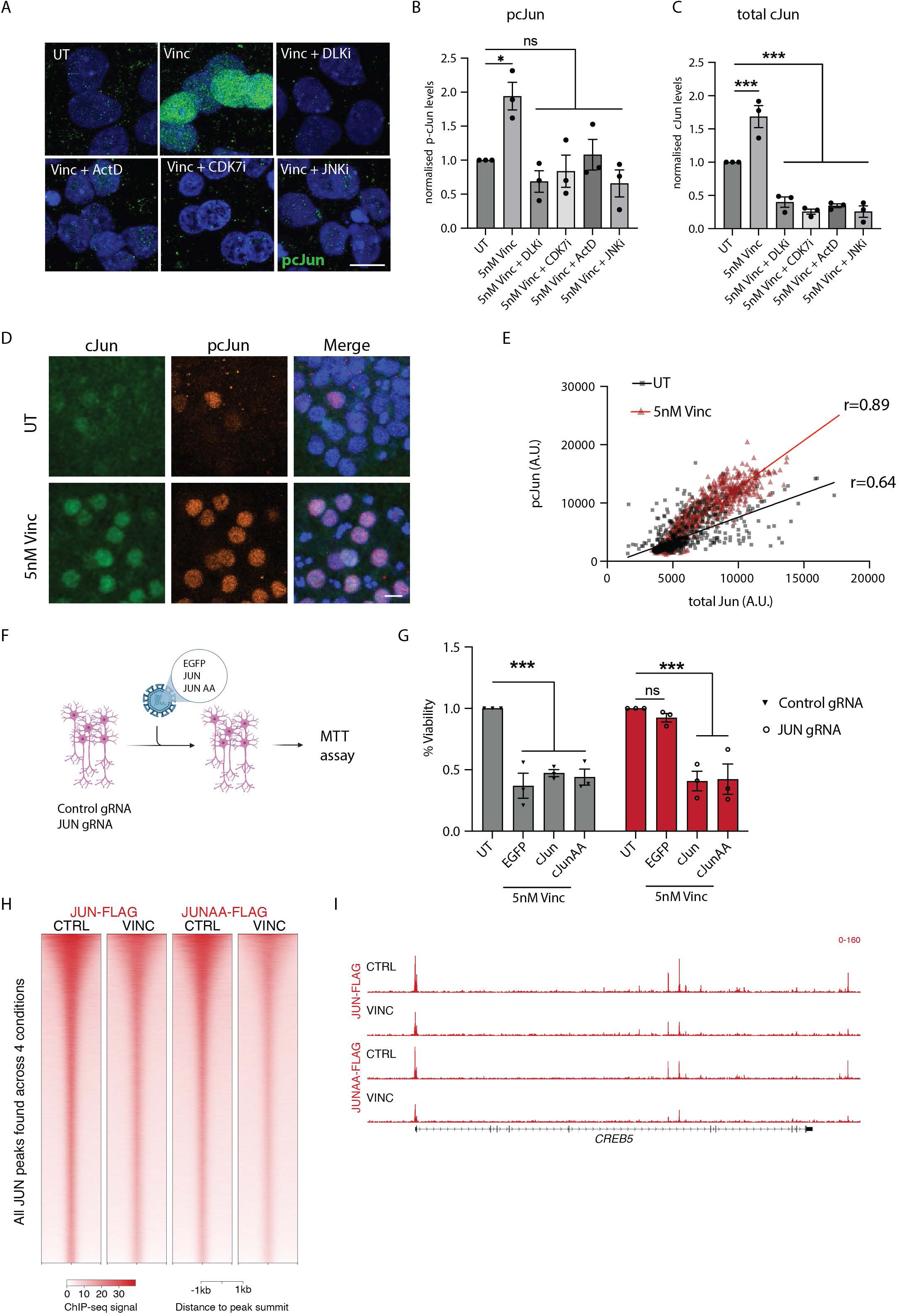
cJun phosphorylation is not required for cell death A. Representative images of cortical i^3^Neurons 24 hours after treatment with 5 nM vincristine, 5 nM vincristine + DLKi (GNE-3511, 500 nM), + JNKi (1 µM), ActD (30 nM), or CDK7i (100 nM). Immunostaining for p-cJun S63 (green), DAPI (blue). Scalebar 15 µm. B. Quantification of relative nuclear p-cJun S63 levels in cortical i^3^Neurons 24 hours after 5 nM vincristine, 5 nM vincristine + DLKi (GNE-3511, 500 nM), + JNKi (1 µM), ActD (30 nM), or CDK7i (100 nM) treatment. Results normalized to UT (untreated). Mean ± SEM. N=3 independent differentiations. One-way ANOVA, Bonferroni correction, ns = not significant, p<0.05 *. C. Quantification of relative nuclear cJun levels in cortical i^3^Neurons 24 hours after 5 nM vincristine, 5 nM vincristine + DLKi (GNE-3511, 500 nM), + JNKi (1 µM), ActD (30 nM), or CDK7i (100 nM) treatment. Results normalized to UT. Mean ± SEM. N=3 independent differentiations. One-way ANOVA, Bonferroni correction, p<0.005 ***. D. Representative images of cortical i^3^Neurons UT and 24 hours after 5 nM vincristine. Immunostaining for cJun (green), p-cJun S63 (orange) and DAPI (blue). Scalebar 15 µm. E. Correlation and Pearson r correlation coefficient of the nuclear levels of p-cJun S63 and total cJun levels in UT (black) and 5 nM vincristine (red) cortical i^3^Neurons for 24 hours. F. Illustration of rescue experiment performed in G. G. Relative viability of control and JUN gRNA cortical i^3^Neurons untreated (UT) or transduced with EGFP, cJun or cJun S63A S73A (JunAA) after 48 hours of 5 nM vincristine. N=3 independent differentiations. Two-way ANOVA, Bonferroni correction, ns = not significant, p<0.005 ***. H. Heatmap showing ChIP-seq signal at all peaks identified across the 4 conditions assessed for cJun. Signal was centered on the summit of the peaks. I. Browser shot showing an example of Jun at a vincristine-responsive transcription factor.

### Vincristine-induced neurotoxicity is driven by cJun but independently of its phosphorylation

We next investigated whether cJun phosphorylation at serines 63 and 73 is necessary to regulate vincristine-induced cell death. We performed a rescue experiment where control and JUN gRNA-expressing neurons were transduced with EGFP, WT cJun or cJun S63A S73A (cJun AA)-expressing lentivirus then treated with vincristine (Fig. 3F). We observed decreased cell viability in control gRNA neurons expressing EGFP, cJun, or cJun AA when exposed to vincristine. JUN gRNA neurons expressing EGFP were protected from vincristine-induced neurodegeneration, but cJun or cJun AA expression rendered them vincristine-sensitive once more (Fig. 3G). This result indicates that phosphorylation of cJun at serines 63 and 73 is not necessary for cJun to drive neuronal cell death in the response to vincristine.

We performed chromatin immunoprecipitation-sequencing (ChIP-seq) to investigate whether cJun phosphorylation alters cJun binding throughout the genome. We lentivirally expressed FLAG-tagged WT cJun and cJun AA in JUN gRNA neurons and treated them with vincristine. JUN KD neurons expressing Jun-FLAG or Jun-AA-FLAG showed a significant decrease in cell viability compared to an EGFP control (Supp. Fig. 4E and F), thereby validating function of the Flag-tagged cJun protein. Although we cannot rule out that overexpression of the Flag-tagged constructs may mask differential binding, ChIP-seq revealed no significant differences in the DNA-binding ability of Jun and Jun AA in DMSO- or and vincristine-treated neurons (Fig. 3H and I). This result shows that cJun is necessary for vincristine-induced neurodegeneration, but that its phosphorylation at serines 63 and 73 is dispensable. It also suggests that other players regulate the transcription of JUN to induce neuronal death. We hypothesized this might be the role of ATF2.

### Vincristine treatment induces p-ATF2 phosphorylation in a DLK/LZK dependent manner

ATF2 and cJun are both AP1 transcription factors^25^, known to form homo- and heterodimers, a feature thought to enhance AP1 family members’ ability to regulate gene expression^25^. ATF2 can be phosphorylated by several kinases including JNK, P38, and ERK^34^ at threonines 69 and 71 (T69 and T71), inducing its translocation to the nucleus ^35^. We hypothesized that vincristine would increase levels of phosphorylated nuclear ATF2 (p-ATF2) in a DLK- and LZK-dependent manner. Treatment of WT but not DLK, LZK or double DLK/LZK KO neurons with vincristine indeed resulted in a significant increase in the levels of nuclear p- ATF2 (Fig. 4A and B). Conversely, treatment of neurons with DLKi and JNKi abolished the increase of nuclear p-ATF2 after vincristine treatment (Fig. 4C). Since ATF2 can be phosphorylated by P38^36^, we verified that levels of phospho-P38 did not increase in neurons treated with vincristine (Supp. Fig. 5A and B). We also confirmed that P38 inhibition did not significantly reduce vincristine-induced neuron death (Supp. Fig. 5C). Together our results show that DLK and LZK regulate ATF2 phosphorylation via the JNK pathway.

**Figure 4:**
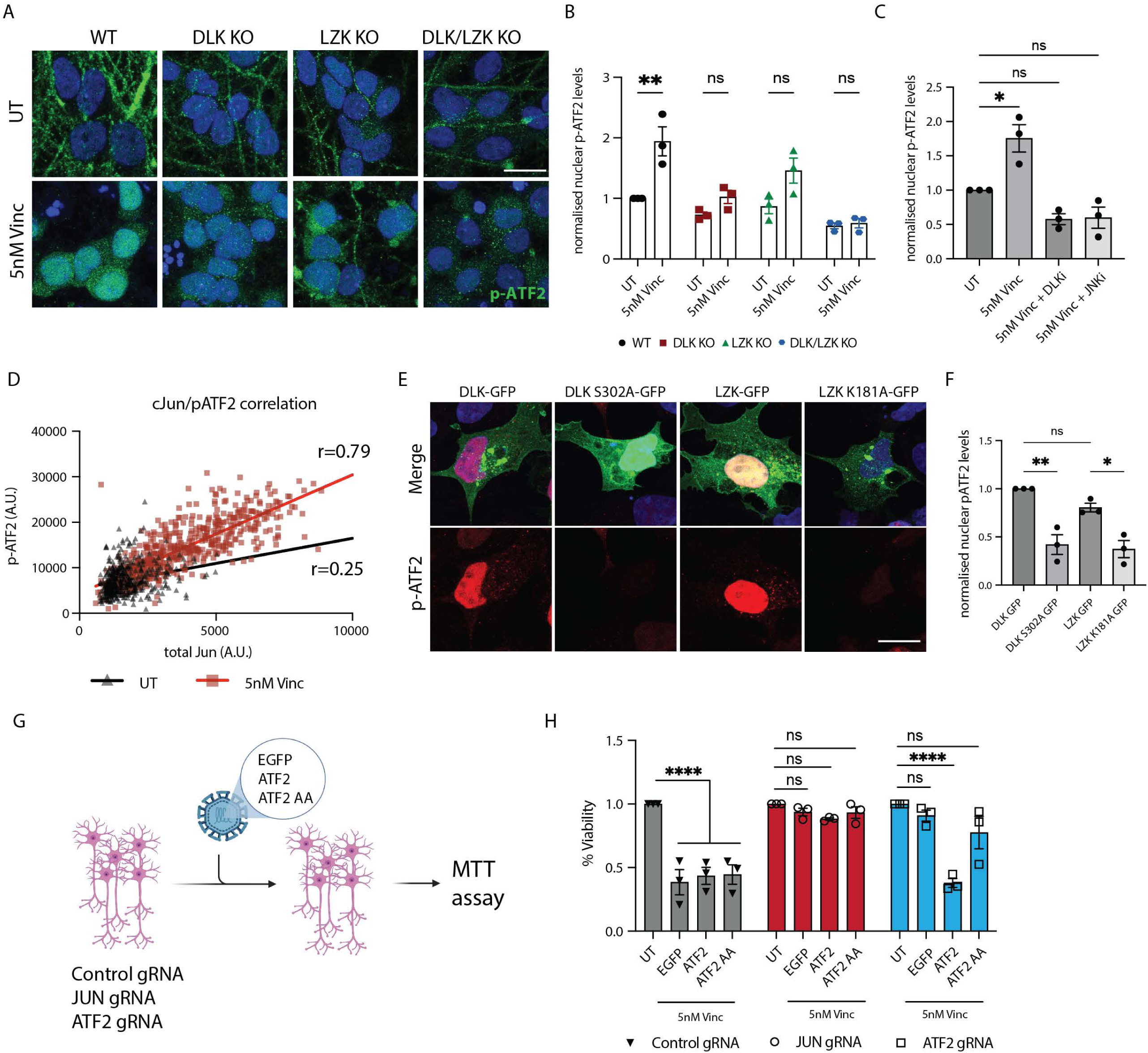
Vincristine increases ATF2 phosphorylation in a DLK/LZK-dependent manner A. Representative images of WT, DLK, LZK and double DLK/LZK KO cortical i^3^Neurons UT and 24 hours after 5 nM vincristine treatment. Immunostaining for p-ATF2 T71 (green), DAPI (blue). Scalebar 20 µm. B. Quantification of relative nuclear p-ATF2 T71 levels UT and after 5 nM vincristine for 24 hours in WT, DLK, LZK and double DLK/LZK KO cortical i^3^Neurons. Results normalized to WT UT. Results represented as mean ± SEM. N=3 independent differentiations. Two-way ANOVA, Bonferroni correction, ns = not significant, p<0.01 **. C. Quantification of relative nuclear p-ATF2 (T71) levels UT and after 5 nM vincristine and 5 nM vincristine + DLKi (GNE-3511, 500 nM), + JNKi (1 µM) for 24 hours in cortical i^3^Neurons Results normalized to UT. Mean ± SEM. N=3 independent differentiations. One-way ANOVA, Bonferroni correction, ns = not significant, p<0.05 *. D. Correlation and Pearson r correlation coefficient of the nuclear levels of p-ATF2 and total cJun levels in UT (Black) and 5nM Vincristine (Red) cortical i^3^Neurons for 24 hours. E. Representative images of HEK-239A cells transfected with DLK-GFP, DLK-S302A- GFP, LZK-GFP and LZK-K181A-GFP for and immunostaining for p-ATF2 (T71) (red), DAPI (blue). (Scale bar = 20 µm). F. Quantification of relative nuclear p-ATF2 (T71) levels in HEK-239A cells transfected with DLK-GFP, DLK-S302A-GFP, LZK-GFP and LZK-K181A-GFP for 24 hours. Results are represented as mean ± SEM. N=3 independent differentiations. One-way ANOVA, Bonferroni correction, ns = not significant, p<0.05 *, p<0.01 **. G. Illustration of rescue experiment performed in H. H. Quantification of the relative viability of control, JUN and ATF2 gRNA cortical i^3^Neurons Untreated (UT) and transduced with EGFP, ATF2 and ATF2 T69A T71A (ATF2 AA) after treatment for 48 hours with 5 nM vincristine. Results are represented as mean ± SEM. N=3 independent differentiations. Two-way ANOVA, Bonferroni correction, ns = not significant, p<0.001 ****.

Moreover, we observed that nuclear p-ATF2 and cJun levels strongly correlated in response to vincristine exposure (red squares, Pearson correlation r=0.79, p<0.0001) and to a lesser extent in untreated neurons (black triangles, Pearson correlation r=0.25, p<0.0001) (Fig. 4D). We further validated that DLK/LZK pathway activation results in increased nuclear p- ATF2 levels in a HEK-293A cell overexpression model. Expression of WT DLK or LZK significantly increases nuclear p-ATF2 levels compared to kinase-dead DLK or LZK (Fig. 4E and F).

### ATF2 phosphorylation is necessary for neurodegeneration

Despite observing a robust increase in the levels in nuclear p-ATF2 after vincristine treatment, our RNAseq data showed that ATF2 RNA levels remained unchanged (Fig. 1H and Supp. Table 1). Phosphorylation of ATF2 at threonine 69 and 71 (T69 and T71) are associated with increased transcriptional activity^25^. We tested whether the phosphorylation of these two residues is necessary for vincristine-induced neuronal death by mutating these two threonines to alanines. Control, JUN-gRNA and ATF2-gRNA neurons were transduced with EGFP, ATF2 or ATF2 T69A T71A (ATF2 AA)-expressing lentivirus then treated with vincristine (Fig 4G). We observed no significant differences in the viability of control neurons expressing any of the three constructs after vincristine treatment, suggesting that overexpression of ATF2 does not exacerbate neuron death. Moreover, knockdown of JUN protected neurons from vincristine-induced cell death when expressing ATF2 and ATF2 AA, suggesting that cJun lies downstream of ATF2. Finally, ATF2 gRNA-expressing neurons transduced with WT but not ATF2 AA degenerated in response to vincristine treatment, indicating that ATF2 phosphorylation is necessary for vincristine-induced neurodegeneration (Fig. 4H).

### ATF2 phosphorylation regulation by DLK following neuron injury is evolutionarily conserved

Next, we asked whether the regulation of ATF2 phosphorylation by DLK and LZK after neuronal injury is evolutionarily conserved, observable *in vivo*, and valid across two different injury paradigms, mild traumatic brain injury (mTBI) and optic nerve crush (ONC).

We have recently shown that after a unilateral mTBI applied to the skull above the motor cortex, ipsilateral cortical neurons in layer V die in a DLK-dependent manner^37^. We hypothesized that mTBI would cause a DLK-driven increase in nuclear levels of pATF2 in layer V neurons. To test this, we deleted DLK in layer V neurons using a layer V-specific Rbp4-Cre driver and a Cre-dependent nuclear reporter (Rbp4-Cre::Dlk^fl/fl^::Sun1-sfGFP). This system allowed us to identify the nuclei of neurons where DLK is deleted. WT and DLK conditional KO (Dlk cKO) mice were subjected to mTBI and harvested 7 days post injury (7 dpi) (Fig. 5A). mTBI produces increased p-ATF2 in the injured neurons of control but not DLK cKO mice (Fig 5B and C). This experiment demonstrates that p-ATF2 increases in neurons after a concussive injury and this increase is DLK-dependent.

**Figure 5:**
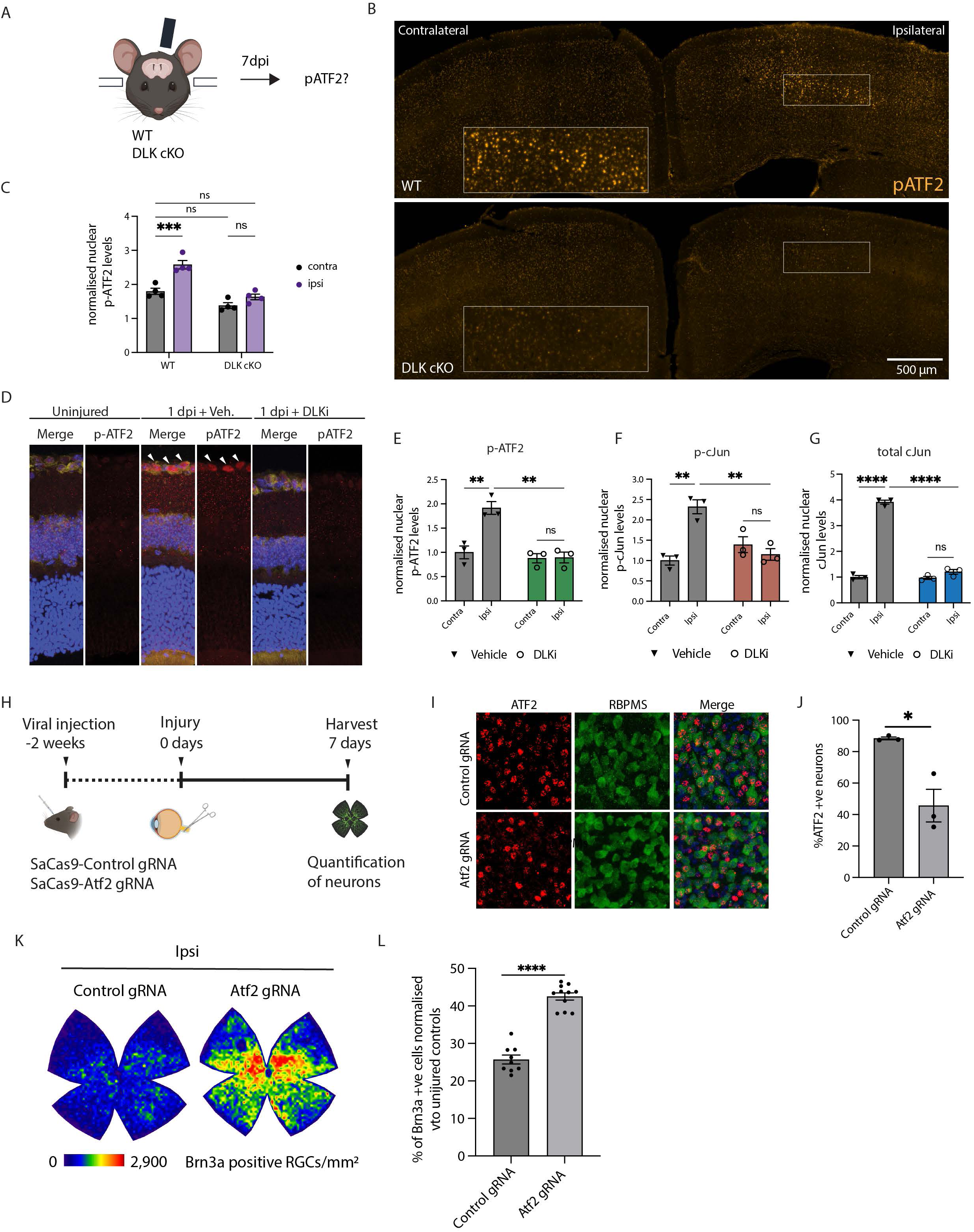
ATF2 phosphorylation is DLK-dependent and ATF2 is required for neuron death in mouse models of neuron injury A. Schematic representation of the experimental design. WT and conditional DLK (DLK cKO) mice were injured. Brains were collected 7 days post injury (7 dpi) and immunostained for pATF2. B. Coronal sections showing pATF2 staining in the injured (ipsilateral) and uninjured (contralateral) cortex in WT (top) and DLK cKO (bottom) mice at 7 days post unilateral mTBI. Insets: enlarged views of boxed areas on the ipsilateral side. C. Quantification of the average pATF2 fluorescence intensity in layer V excitatory neurons revealing a DLK-dependent increase in ATF2 phosphorylation in ipsilateral neurons at 7 dpi. Results represented as mean ± SEM. Each dot represents one animal and the average of all the Cre+ nuclei in 3-4 ROIs per section across 3-4 sections per animal. Two-way ANOVA, Bonferroni correction, ns = not significant, p<0.001 ***. D. Representative images of retinas uninjured and 1 dpi in mice treated with vehicle (Veh.) or DLK inhibitor GNE-3511 (DLKi). Immunostained with p-ATF2 T71 (red), neurotrace (yellow) and DAPI (blue). E. Quantification of relative nuclear p-ATF2 T71 levels in the nucleus of retinal ganglion cells (RGCs) in mice treated with vehicle or DLK inhibitor (DLKi) contralateral (contra) and ipsilateral (ipsi) 1 day after optic nerve crush. Results represented as mean ± SEM. N=3 mice. Two-way ANOVA, Bonferroni correction, ns = not significant, p<0.01 **. F. Quantification of relative nuclear p-cJun S63 levels in the nucleus of RGCs in mice treated with vehicle or DLK inhibitor GNE-3511 at 1 dpi. Mean ± SEM. N=3 mice. Two- way ANOVA, Bonferroni correction, ns = not significant, p<0.01 **. G. Quantification of relative total cJun levels in the nucleus of RGCs in mice treated with vehicle or GNE-3511 at 1 dpi. Mean ± SEM. N=3 mice. Two-way ANOVA, Bonferroni correction ns = not significant, p<0.001 ****. H. Schematic representation of RGC survival following optic nerve crush (ONC). I. Representative images of control and ATF2 gRNA retinas 3 weeks after gRNA transduction. Immunostained with ATF2 (red) and RBPMS (green). J. Quantification of ATF2 positive (+ve) neurons control and ATF2 gRNA retinas 3 weeks after gRNA transduction. N = 3 mice per condition. Unpaired t test, p < 0.05 *. K. Representative isodensity maps display the topological survival of Brn3a+RGCs at 7 dpi. ATF2 knockdown delays ONC-induced RGC degeneration across the retina. Colorscale for isodensity maps ranges from 0 (purple) to 2900 (red) RGCs/mm^2^. L. Quantification of percentage Brn3a-positive RGCs in retinas of control and ATF2 gRNA mice at 7 dpi (normalized to contralateral condition). N = 9-11 mice per condition. Unpaired two-tailed t test, p < 0.0001 ****.

The second model we used, optic nerve crush (ONC), induces transcription-dependent neuronal death regulated by DLK^38–40^. Mice treated with vehicle or GNE-3511 (DLKi) were subjected to an ONC and harvested one day post injury (1 dpi). We evaluated the presence of p- ATF2 in the nuclei of retinal ganglion cells (RGCs) by immunofluorescence. We observed pATF2 significantly increased with p-cJun and total cJun levels in vehicle-treated ipsilateral RGCs but not in those treated with DLKi (Fig. 5D-G), again demonstrating that p-ATF2 increases in injured neurons in a DLK-dependent manner.

Taken together, our results show that the phosphorylation of ATF2 in response to neuronal stress is driven by the DLK/LZK pathway both *in vivo* and *in vitro*, and across chemical, mechanical and traumatic injury paradigms.

### ATF2 regulates neuronal death after axon injury *in vivo*

Since we observed that phospho-ATF2 is upregulated in retinal ganglion cells after ONC, we asked whether ATF2 knockdown is neuroprotective in this injury model. We transduced RGC neurons with Cas9 from Staphylococcus aureus (SaCas9) and control gRNA or gRNA targeting mouse ATF2. After 2 weeks to allow for viral expression and ATF2 knockdown, we performed optic nerve crush injuries. One week later (7 dpi), we harvested the retinas, immunolabeled them for markers of RGCs and counted surviving neurons (Fig. 5H). We used uninjured retinas to assess the extent of ATF2 reduction by the virally delivered Cas9 and gRNA. We observed that knockdown of ATF2 did not result in loss of RGCs in uninjured retinas (Supp. Fig 6B and C), but that though ATF2 expression was reduced, it was incompletely knocked down (Fig. 5K and L and Supp Fig. 6D). Importantly, despite the incomplete knockdown, we found that ATF2 knockdown significantly improved RGC survival following ONC (Fig. 5H and I; control gRNA: 25.7 ±1.2 vs ATF2 gRNA: 42.3± 1.0 (average ± SEM)). The partial knockdown of ATF2 we observed suggests that a more thorough reduction could result in even greater protection.

**Figure 6:**
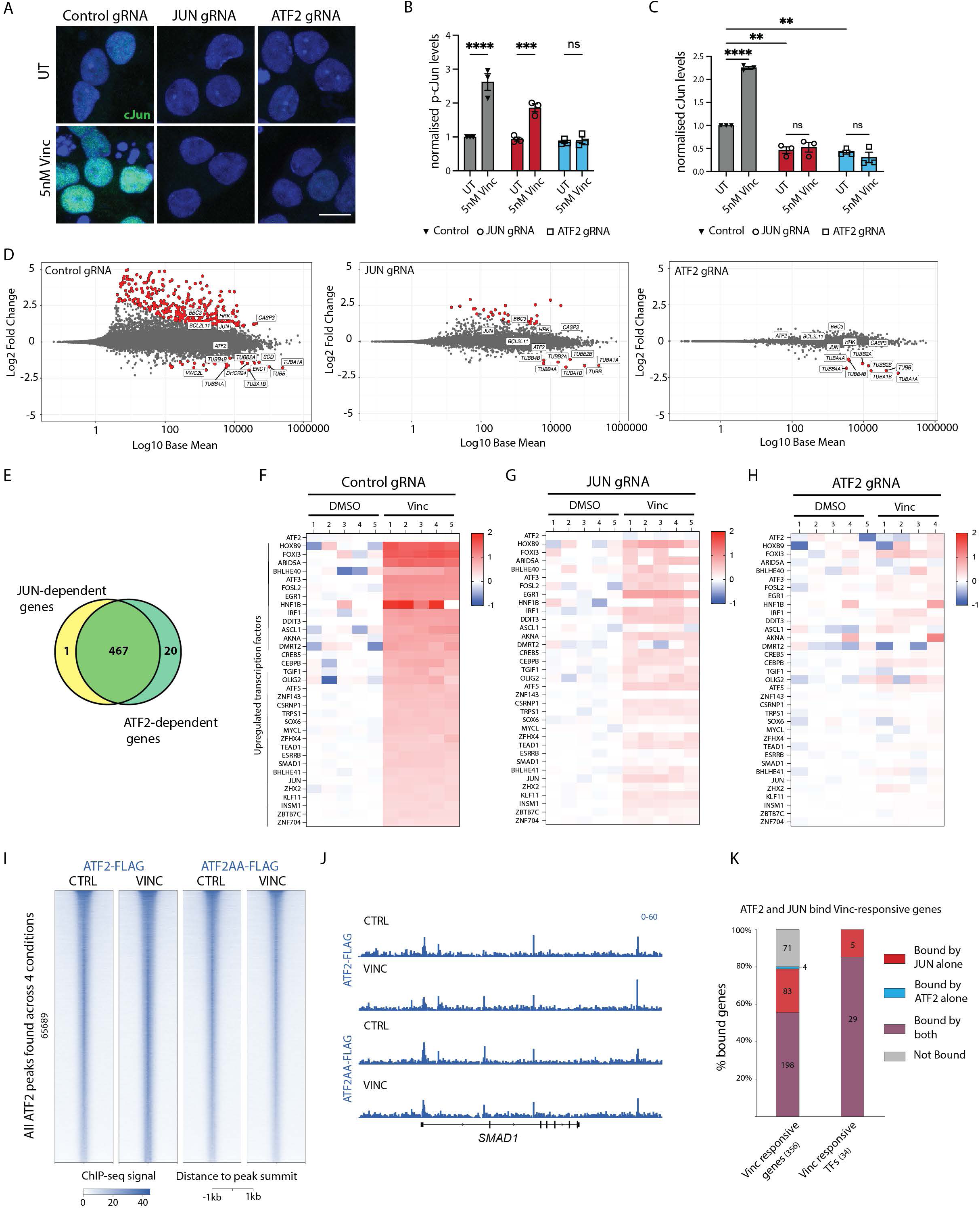
ATF2 regulates JUN transcription A. Representative images of control, JUN gRNA and ATF2 gRNA cortical i^3^Neurons 24 hours after 5 nM vincristine treatment. Immunostaining for total cJun (green), DAPI (blue). Scalebar 20 µm. B. Quantification of relative nuclear p-cJun S63 levels in control, JUN gRNA and ATF2 gRNA cortical i^3^Neurons 24 hours after 5 nM vincristine treatment. Results normalized to control gRNA UT (untreated). Mean ± SEM. N=3 independent differentiations. One- way ANOVA, Bonferroni correction, ns: not significant, p<0.005 ***, p<0.001 ****. C. Quantification of relative nuclear total cJun levels in control, JUN gRNA and ATF2 gRNA cortical i^3^Neurons 24 hours after 5 nM vincristine. Results normalized to control gRNA UT. Mean ± SEM. N=3 independent differentiations. One-way ANOVA, Bonferroni correction, ns = not significant, p<0.01 **, p<0.001 ****. D. MA plots showing the transcriptional changes in control, JUN and ATF2 gRNA cortical i^3^Neurons 16 hours after 5 nM vincristine. Significantly differentially expressed genes labelled in red. E. Venn diagram showing common genes regulated by DLK and LZK after vincristine treatment. F. Heatmap of normalized levels of transcription factors that are significantly upregulated in control gRNA cortical i^3^Neurons 16 hours after 5nM vincristine. Results normalized to control gRNA UT. Each column represents a biological replicate. G. Heatmap of normalized levels of transcription factors that are significantly upregulated in JUN gRNA cortical i^3^Neurons 16 hours after 5 nM vincristine. Results normalized to JUN gRNA UT. Each column represents a biological replicate. H. Heatmap of normalized levels of transcription factors that are significantly upregulated in ATF2 gRNA cortical i^3^Neurons 16 hours after 5 nM vincristine. Results normalized to ATF2 gRNA UT. Each column represents a biological replicate. I. Heatmap showing ChIP-seq signal at all peaks identified across the 4 conditions assessed for ATF2. Signal was centered on the summit of the peaks. J. Browser shot showing that ATF2 binds strongly to the SMAD1 promoter. K. Number of vincristine-responsive genes or transcription factors that are bound by cJun, ATF2, both or none.

### ATF2 regulates cJun and p-cJun levels after vincristine treatment

Using luciferase reporters, it has been shown that ATF2 and cJun form heterodimers that induce transcription by binding to the AP-1 site of the JUN promoter^41^. We asked whether ATF2 regulates total cJun and phospho-cJun levels after injury. We found that after vincristine treatment, ATF2 knockdown prevented cJun phosphorylation (Fig. 6A and B). Total cJun levels were increased in control but not cJun or ATF2 KD neurons (Fig. 6C). These results identify ATF2 as a key regulator of both total cJun protein levels and its phosphorylation.

### ATF2 and cJun regulate a common set of genes in response to vincristine

To characterize whether ATF2 and cJun regulate a common set of genes, we performed RNAseq on control, JUN and ATF2 gRNA-transduced neurons treated vincristine. *ATF2* and *JUN* transcripts were significantly reduced in ATF2 and JUN gRNA neurons, validating our knockdown approach (Supp. Fig. 7A and B). Consistent with the observed decrease in total cJun protein levels in untreated ATF2 gRNA neurons (Fig. 6C and Supp. Table 3), *JUN* transcripts were also significantly downregulated (Supp. Fig. 7A and Supp. Table 3) indicating that under homeostatic conditions, ATF2 regulates cJun protein levels at the transcriptional level.

As expected, control neurons underwent a robust transcriptional response to vincristine exposure, and pro-apoptotic genes such as *CASP3*, *BBC3* (which encodes PUMA), *JUN*, *BCL2L11* and *HRK* were significantly upregulated (but not *ATF2*, which is not transcriptionally regulated). The expression of all these genes was attenuated in both JUN and ATF2 KD neurons (Fig. 6D). Genes regulated by JUN and ATF2 overlapped almost completely (Fig. 6E). As a result, we conclude that ATF2 and cJun regulate neuron degeneration within the same common pathway.

We had previously observed that ATF2 is the only transcription factor hit from our CRISPRi screen that is not transcriptionally upregulated upon vincristine treatment (Fig 2E). We now asked whether ATF2 acts as a master transcription regulator of transcription factors in response to vincristine. We find that RNA levels of 34 transcription factors, but not ATF2, are significantly upregulated in control neurons in response to vincristine (Fig. 6F). Levels of these TFs are partially reduced in JUN knockdown neurons (Fig. 6G), but completely suppressed in ATF2 knockdown neurons (Fig. 6H), suggesting that ATF2 is indeed the core transcription factor controlling this program.

### ATF2 phosphorylation does not regulate its DNA binding ability

We next performed ChIP-seq to ask whether ATF2 phosphorylation affects its ability to bind DNA. For this we expressed N-terminally FLAG-tagged WT ATF2 or ATF2 T69A T71A (ATF2-AA) in ATF2 knockdown neurons for 2 days. We did this in neurons treated with vincristine for 24 hours or DMSO as a control. We validated the ability of both FLAG-tagged constructs to signal by expressing them in ATF2 knockdown neurons. We found that FLAG-

ATF2 but not FLAG-ATF2-AA is able to rescue the decreased levels of cJun protein observed in ATF2 KD neurons (Supp. Fig. 7C and D). Surprisingly, our ChIP-seq results revealed no significant differences in the DNA-binding ability of ATF2 and ATF2 AA in control and vincristine-treated neurons (Fig. 6I and J).

Lastly, we asked whether cJun and ATF2 work together or independently to drive the pro-degenerative transcription cascade. We combined our ChIP-seq results and found that as expected, at many regions, JUN and ATF2 showed similar enrichment (Supp. Fig. 7E middle cluster). Importantly, even though we used an ectopic expression system, we also identified many regions with differential enrichment, showing that ATF2 and JUN do not completely overlap in their genomic targets (Supp. Fig. 7E top cluster shows regions more strongly bound by ATF2 and bottom cluster regions preferentially bound by JUN and Supp. Fig.7F). While we noticed that regions with more ATF2 enrichment were also partially bound by JUN, the reverse was not true. Most regions with higher JUN binding showed almost no ATF2 enrichment. These data suggest that JUN-ATF2 and JUN-other TFs bind as heterodimers on the common sites, but JUN binds with other partners at the non-ATF2 sites.

We analyzed whether ATF2 and cJun bind to vincristine-induced upregulated genes. We found that a large proportion of the upregulated genes (198/356 or 55.6%) were bound by both ATF2 and cJun (Fig. 6K). Interestingly, the majority of upregulated transcription factors in response to vincristine were bound by both ATF2 and cJun (29/34 or 85%) (Fig. 6K).

Together our results identify ATF2 as an essential relay between the DLK-LZK-JNK signaling cascade and transcription. ATF2 phosphorylation results in its translocation to the nucleus and is a key effector that initiates a pro-degenerative transcriptional program in which an essential initial step is upregulation of JUN.

## Discussion

Here, we demonstrate that DLK and LZK both regulate neuronal death through phosphorylation of ATF2 *in vivo* and *in vitro*. Under conditions of neuronal stress, levels of phosphorylated ATF2 rise within the nucleus, promoting the transcription of *JUN*. We establish that, although cJun is essential for neuronal degeneration, this process is independent of its phosphorylation. Consequently, our findings unveil ATF2 as an evolutionarily conserved and critical component of the DLK/LZK signaling pathway and a novel potential therapeutic target for neurodegenerative disorders in which the DLK pathway has been identified to play a role such as Alzheimer’s disease^8,42^ and ALS^42,43^. We find that ATF2 acts as the key transcriptional switch that converts the DLK/LZK phosphorylation cascade into a transcriptional response.

In mice, phosphorylation of cJun and neuronal death following NGF withdrawal and other models of axonal injury are almost completely driven by DLK^33,38,44,45^. However, in human neurons, we find that these processes are partially driven by DLK and LZK individually, with both proteins being necessary to cooperatively induce neurodegeneration. We saw no differences in the levels of p-ATF2 induced by DLK and LZK overexpression in HEK-293A cells, suggesting that DLK and LZK are equally potent kinases. We also show that DLK and LZK regulate a common set of genes. Aside from several tubulin-encoding genes, the transcriptional response to vincristine is entirely dependent on both DLK and LZK. We therefore identify LZK together with DLK as master regulators of neurodegeneration in human neurons.

We demonstrate that the transcriptional regulation exerted by cJun under cytotoxic stress is independent of its phosphorylation. The increase in phospho-cJun levels is only a correlate of an increase in total cJun RNA and protein, which are critical for the JUN-dependent transcriptional response and neuronal death whereas the phosphorylation is not. Upon close examination of the literature, we found that a standard practice in the field has been to report a stress-induced increase in cJun phosphorylation by normalizing phospho-cJun levels to the loading control, rather than to total Jun protein levels. The question of whether the phosphorylation of cJun is necessary for neurodegeneration has been highly debated^14–18^, however our results indicate that in human neurons, cJun is actively transcribed under stress conditions and drives neurodegeneration regardless of its phosphorylation status.

Here we present the first whole genome CRISPR interference (CRISPRi) screen to identify modifiers of chemotherapy-induced neurotoxicity in human neurons. We identify ATF2 as a key player and elucidate its lesser-known role as a driver of neurodegeneration. Studies in stable cell lines have shown that localization of ATF2 to the nucleus is dependent on its phosphorylation at residues T69 and T71^35^. While elevated nuclear ATF2 levels have been reported in AD^46,47^, Parkinson’s disease models^48^ and stroke^49^, little is known about the mechanisms driving its translocation to the nucleus or its downstream targets in neurons. Here we identify DLK and LZK as novel regulators of ATF2 phosphorylation and nuclear localization *in vivo* and *in vitro*. It has been suggested that cJun and ATF2 belong to different branches of the DLK/LZK pathway, and that simultaneous silencing of both factors provides an enhanced neuroprotection^4^. We find that ATF2 regulates cJun transcription as well as total and phospho-cJun levels after stress, that p-ATF2 and cJun levels strongly correlate in nuclei of stressed neurons, and that ATF2 and cJun control a highly overlapping set of genes. We therefore conclude that ATF2 and cJun belong to a unified signaling pathway where ATF2 resides upstream of cJun and downstream of DLK/LZK and JNK (Fig. 7).

**Figure 7:**
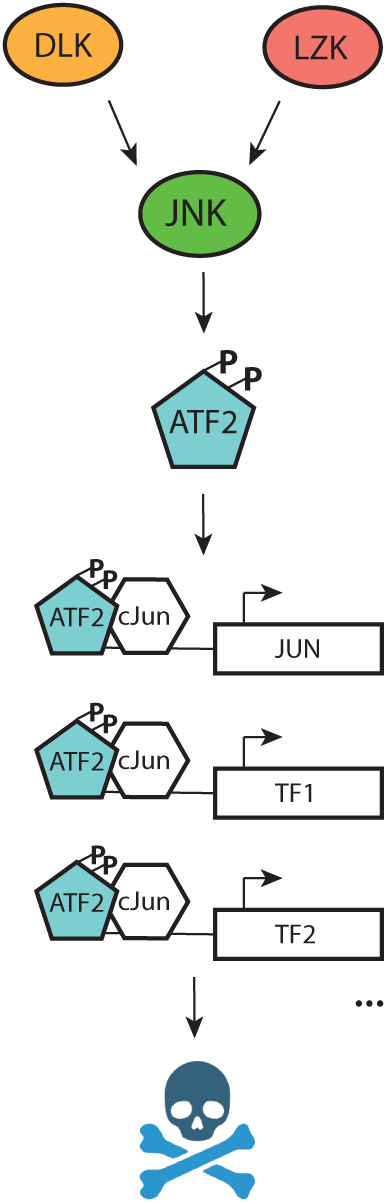
Schematic representation of the proposed DLK/LZK signaling pathway

Our work also opens the door to systematically identify the downstream targets of transcription factors in health and disease in human neurons. The generation of virtually pure human neuronal subtypes eliminates transcriptional contamination from other cell types and reduces the costs associated with addressing these questions. Other transcription factors, such as ATF4^23^, CHOP^21^ and CREB^50^ have been implicated in neurodegeneration. Indeed, a recent report described that a core set of transcription factors, ATF3, DDIT3, CEBPG and ATF4 (but not ATF2) collaborate to regulate the chromatin accessibility in neurons in response to injury^51^. Interestingly, we find that two of these transcription factors (ATF3 and DDIT3) are upregulated in neurons in an ATF2- dependent manner. Elucidating how these and other transcription factors signal, as well as exploring the overlap between them, will provide an invaluable tool for the neurodegenerative research community.

We have identified ATF2 as an evolutionarily conserved component of the DLK/LZK pathway. Although silencing ATF2 itself may prove therapeutically challenging, targeting its pathological interactions with upstream and downstream effectors may be exploited as a potential therapeutic target. Critical questions should therefore be addressed in future, such as how is ATF2 imported into the nucleus in human neurons? How does the phosphorylation of ATF2 affect its interaction with other proteins? Can the interaction of JNK and ATF2 can be blocked? This work also identifies multiple drivers of neurodegeneration that should be further explored by the field and highlights the feasibility of using small molecule cellular stressors to perform large scale survival screens to identify disease-relevant candidate genes.

## Methods

### Immunofluorescence

Immunofluorescence was performed as previously described^52^. Briefly, 10-14 day old neurons were treated with the appropriate stressor and inhibitors and fixed with ice cold 4% PFA for 10 minutes at room temperature. Cells were washed once with PBS and permeabilized with 0.1% TritonX in PBS for 5 mins and blocked in 5% normal donkey serum (NDS) in PBS at room temperature for 1 hour. Cells were stained with the selected primary antibody overnight at 4 degrees in 2.5% Donkey serum. Cells were washed 3 times in PBS and stained with the appropriate secondary antibody at a concentration of 1:500 in 2.5% NDS for 1 hour at room temperature. Cells were washed three times with PBS and DAPI was used as a nuclear counterstain. Antibodies used: p-cJun (S63) (CST, #9261S), total cJun (CST, #9165), p-ATF2 (T71) (Abcam, #ab32019), 488-conjugated cJun (SCBT, #sc-74543 AF488), pP38 (Y180, Y182) (CST, #9211).

### Western blotting

Western blotting was performed as previously described^52^. Briefly, cells were harvested in ice cold RIPA buffer (Invitrogen, #89901) supplemented with protease inhibitors (Sigma, 11836153001) and PhosStop (Sigma, #4906845001). Protein concentrations were calculated using a BCA assay (ThermoFisher, #23225). 10-20 µg of total protein lysate was loaded into a 4–20% Mini-PROTEAN® TGX™ Precast Protein Gels (Biorad, #4561095). Gels were transferred into PVDF membranes and blocked for 1 hour at room temperature in 5% BSA. Membranes were incubated with the primary antibody (primary antibodies, catalog numbers and concentrations listed below) in 2.5% BSA solution in a cold room, rocking overnight. Membranes were washed 3 times with PBS 0.1% Tween solution. Secondary antibody (goat anti-rabbit HRP and IRDye^®^ 680RD Goat anti-Mouse IgG) was incubated for 1 hour at room temperature at a concentration of 1:8000 in 2.5% BSA solution. Membranes were washed 3 times with PBS 0.1% Tween solution prior to developing. Western blots were developed using Clarity Western ECL Substrate (Biorad, #1705061) or fluorescence and imaged using a ChemiDoc MP Imaging system (Biorad). Band intensity was quantified using FIJI imaging analysis software. Antibodies used: γH2AX (CST, #9261S), DLK (Sigma #HPA039936), LZK (Sigma, #HPA016497), b Actin (Sigma, # A1978, 1:5000), GAPDH (CST, #97166S).

### Sensory neuron differentiation

Sensory neurons were differentiated as previously described^28^. Briefly, i^3^ iPSCs were dissociated using Accutase (Life Technologies, #A1110501). Cells were plated in Matrigel-coated (1:100 Corning) plates in sensory neuronal induction media on day 0 (1:1 mix of Neurobasal (ThermoFisher #21103049) and DMEM/F12 medium (ThermoFisher #11320-033) supplemented with1× MEM nonessential amino acids (NEAA) (ThermoFisher, #11140050), 10 μM S1049 inhibitor (Y-27632; Tocris, #1254), 2 μg/ml doxycycline (Clontech, #631311), 1X N2 supplement (Life Technologies, #17502048), 1× B27 Plus Supplement (ThermoFisher Scientific, #A3582801). Induction media was changed daily for 2 days. On day 3 of differentiation, cells were dissociated using Accutase and plated in dishes coated with Polyethyleneimine (Sigma, #408727-100ML) and laminin (15ng/ml Corning, #354232). Cells were plated in sensory neuronal maturation media (1:1 mix of Neurobasal (ThermoFisher #21103049) and DMEM/F12 medium supplemented with1× MEM nonessential amino acids (NEAA) (ThermoFisher, #11140050), 2 μg/ml doxycycline (Clontech, #631311), 1X N2 supplement (Life Technologies, #17502048), 1× B27 Plus Supplement (ThermoFisher Scientific, #A3582801), 10 ng/ml BDNF (PeproTech, #450-02), 10 ng/ml NT-3 (PeproTech, #450-03), 10 ng/ml GDNF (PeproTech, #450-10 – 100 μg), 10 ng/ml NGF (R&D systems, #256-GF-100), 1 mg/ml mouse laminin (Invitrogen, #23017015). Half of the neuronal maturation media was removed and replenished with fresh media every 2-3 days.

### Cortical neuron differentiation

I^3^Neurons were differentiated as previously described^52^. Briefly, i^3^ iPSCs were dissociated using Accutase (Life Technologies, #A1110501). Cells were plated in Matrigel-coated (1:100 Corning) plates in neuronal induction media on day 0 (Knockout Dulbecco’s modified Eagle’s medium (DMEM)/F12 medium; Life Technologies Corporation, #12660012), 1X N2 supplement (Life Technologies, #17502048), 1× GlutaMAX (ThermoFisher Scientific, #35050061), 1× MEM nonessential amino acids (NEAA) (ThermoFisher Scientific, #11140050), 10 μM ROCK inhibitor (Y-27632; Tocris, #1254), and 2 μg/ml doxycycline (Clontech, #631311). Neuronal induction media was changed once a day for 2 more days. On day 3 of induction, cells were dissociated using Accutase and plated in dishes coated with poly-L-ornithine (PLO; 0.1 mg/ml; Sigma, #P3655-10MG). Cells were plated in neuronal maturation media (BrainPhys medium (STEMCELL Technologies, #05790), 1× B27 Plus Supplement (ThermoFisher Scientific, #A3582801), 10 ng/ml BDNF (PeproTech, #450-02), 10 ng/ml NT-3 (PeproTech, #450-03), 1 mg/ml mouse laminin (Invitrogen, #23017015), and 2 μg/ml doxycycline). Half of the neuronal maturation media was removed and replenished with fresh media every 2-3 days.

### Generation of DLK and LZK KO iPSCs

DLK and LZK KO neurons were generated as previously described^53^. WT cortical and sensory iPSCs were transfected with two gRNAs targeting exon 2 of the DLK gene gRNA1: CCCAGGCTCCCTGCTACTGCAT and gRNA2: TCCTTTGGCGTGGTGCTATGGG or LZK gene gRNA1: TTGAGAAGATCAAAAGTGAAGG and gRNA2: AGATGAAGAACTGATTCGAAGG and a Cas9-GFP expressing constructs. iPSCs were allowed to grow for two days, then colonies were dissociated to make single cells using Accutase for 10 minutes at 37°C and a serial dilution was performed in two 6-well plates in mTESR Plus Basal Media with ROCK inhibitor. iPSCs were allowed to grow until single colonies were visible, hand-picked using a P1000 pipette, and an individual colony was transferred to a well of a 24-well plate. Once confluent, 10% of the cells were used for passaging and DNA was extracted on the remaining 90% of the cells using a Qiagen DNA extraction kit (Qiagen; #69504). Clones were screened by PCR (DLK KO primers: Forward: TCAGGTGAATGCTGAGCCAGC Reverse: TGGAGACTGTTGCTTCCCACACLZK KO primers: Forward: TCAGGGATTGAAACTCCTGGAA Reverse: GGGTGCCTGTTAACATCTGAAG). Cortical double DLK LZK KO cells were generated by additionally knocking down DLK in LZK KO iPSCs as described above. Sensory double DLK LZK KO cells were generated by additionally knocking down LZK in DLK KO iPSCs as described above. Single and double KO cortical and sensory neurons were further validated by Western blotting.

### CRISPRi library lentiviral packaging

The Human Genome-Wide CRISPRi Dual-sgRNA Library (Cellecta KIDHGW-105K-P, 5 CRISPRi dual-sgRNA construct/gene, subsequently referred to as KIDHGW) was packaged as follows: 80% confluent Lenti-X™ 293T cells (Takara Bio; #632180) were transfected in 15-cm plates using Lipofectamine 3000 transfection reagent (ThermoFisher; #L3000015) with 19.5 μg library plasmid, 13.3 μg psPAX2, 4.5 μg pMD2.G (Gift from Didier Trono; Addgene #12260 and 12259), 1.8 μg pAdvantage (Promega; #E1711) and 80 μL P3000 reagent diluted in 2.4 μL Opti-MEM I reduced serum medium (Thermofisher #31985070) and 60 µL Lipofectamine 3000 transfection reagent diluted into another 2.4 mL Opti-MEM and incubated at room temperature for 5 min. Diluted Lipofectamine 3000 was added to DNA mixture and incubated for 10 min and added dropwise to plates. Sixteen hours later, medium was replaced with DMEM 10% FBS with 1x ViralBoost (Alstem; #VB100). Two days later, medium was collected, centrifuged and the supernatant was filtered through a 0.45um filter, mixed well with Lentivirus Precipitation Solution (Alstem; #VC100), incubated at 4°C overnight followed by centrifugation according to the manufacturer’ protocol. Viral pellet was resuspended in Essential 8™ Flex Medium (ThermoFisher; #A2858501) in 1/40^th^ of the collected media volume, aliquoted and stored at - 80°C. Library titration was performed by infection of IPSCs^ZIM3-dCas9^ with different virus dilutions followed by RFP assessment by flow cytometry on day 3 post infection.

### Vincristine CRISPRi positive selection screen

IPSCs^ZIM3-dCas9^ were dissociated by StemPro™ Accutase™ Cell Dissociation Reagent (ThermoFisher; #A1110501), counted and resuspended in Essential 8™ Flex Medium supplemented with 10 μM Y-27632 (Tocris; #1254) and 8 μg/ml polybrene (Thomas Scientific; #C788D57) and mixed with a volume of KIDHGW lentivirus corresponding to an 1600x representation with MOI of 0.3. Cells were then seeded on Matrigel-coated 15 cm dishes (20 million cells per 20 ml of Essential 8™ Flex Medium supplemented with 10 μM Y27632 and 8 μg/ml polybrene per 15 cm dish) overnight, followed by fresh medium replacement. One day later, medium was replaced with fresh Essential 8™ Flex Medium supplemented with 0.5 µg/mL puromycin (ThermoFisher; #A1113803) for 3 days. Selected cells were dissociated and seeded on new Matrigel-coated 15 cm dishes and taken through the 3 days pre-differentiation followed by seeding on poly-ornithine-coated dishes for cortical neuronal differentiation for 7-8 days while maintaining >1600x representation with 12 million cells per 15 cm dish in 42 plates.

Terminally differentiated cortical neurons ^ZIM3--dCas9^ were split into 2 groups: 1-treated with 10 nM vincristine (21 plates) or 2-DMSO vehicle (21 plates) for 48 hrs then dissociated following a protocol adapted from Kirwan *et. al.* ^54^. We selected 10 nM vincristine because we found that at this concentration 100% of the cells degenerate. Briefly, cells were washed in PBS and dissociated using TrypLE (ThermoFisher; #12604021) with 5 units/ml papain (Worthington; #LK003178) and 1mM EDTA, resuspended in BrainPhys™ Neuronal Medium (Stem Cell Technologies; #05790) supplemented with DNAse I (1000 units/30 ml) (Worthington; #LK003172), 10 μM Y-27632 and 0.5 mM EDTA, pelleted, washed, re-pelleted and snap frozen.

Genomic DNA was harvested with NucleoSpin Blood XL, Maxi kit for DNA from blood (Macherey-Nagel) or QIAamp® DNA Mini (Qiagen) including proteinase K digestion, for control and vincristine-treated groups, respectively. The screen was repeated twice with the second screen having an additional 10 min incubation with DNase I (Worthington Biochemical Corporation) at 37 °C prior to dissociation. DNA was amplified and sequenced by Cellecta with >500 reads/sgRNA according to their NGS prep kit (#LNGS-400) and sequenced on a NextSeq 500 (Illumina). Counts files of DMSO and vincristine -treated samples subject to comparison were input into MAGeCK-including Negative Controls (https://kampmannlab.ucsf.edu/mageck-inc) where log_2_ fold changes (LFCs) and p values were calculated for each sgRNA using the ‘mageck test –k’ command, followed by Robust Rank Aggregation (RRA). RRA fits a negative binomial model to test whether the initial sgRNA counts (DMSO) vary significantly from the final condition’s counts (Vincristine treatment)^55^. The resulting guide-level p values are combined at the gene level using a modified robust ranking algorithm, with corresponding estimated false discovery rates (FDRs) in the gain-of-function screen (vincristine resistance).

### MTT assay

After differentiation, 50,000 cells per well were plated in 96 well plates (Thomas Scientific, #229196) and differentiated for 10-14 days. Cells were treated with the appropriate stressor in 100 µl of neuronal medium. After treatment, MTT assay (Sigma, #CT02) was performed following the manufacturer’s recommendations. Briefly, 10 µL of solution A was added to each well and placed in the incubator for 4 hours. 100 µL of solution B was added to each well and placed in the incubator overnight. Cell viability was measured using a microplate reader at 570nm. Chemical inhibitors and concentrations used: Actinomycin D (ActD) (Selleckchem, #S8964; 3 nM), THZ1 2HCL (CDK7i) (Selleckchem, #S7549; 100 nM), DLKi (GNE-3511) (Cayman Chemical, #19174; 500nM), Vincristine (Sigma, #V8879-5MG), P38 inhibitor (SB202190) (Tocris, #1264; 100 nM), JNKi (SP600125 1µM, Selleckchem, #S1460).

### RNA extraction for RNA seq

For RNA extraction, 2.5 million neurons were pleated in 6 well plates (Thomas Scientific, #1198D71) and differentiated for 10-14 days. Neurons were treated with DMSO or 5 nM vincristine for 16 hours before RNA extraction. Neurons were washed once with PBS and harvested in 700 µL Qiazol (Qiagen, #79306) and immediately placed on ice. RNA extraction was performed using RNeasy Mini Kit (Qiagen, #74104) following the manufacturer’s instructions. Columns were treated with RNase-Free DNase (Qiagen, #79254), and RNA was eluted in 25 µL of RNAse free water. Eluted RNA was separated into two tubes, one containing 2 µL, subsequently used for quality control and the other used for RNAseq.

For RNA sequencing, 1-4 µg of total RNA samples were purified with PolyA extraction, and then purified mRNAs were constructed to RNA-Seq libraries with specific barcodes using Illumina TruSeq Stranded mRNA Library Prep Kit. All the RNA-Seq libraries were pooled together and sequenced using Illumina NovaSeq 6000 to generate approximately ∼40 million 2x100 paired-end reads for each sample. The raw data were demultiplexed and analyzed further.

### RNA seq analysis

Sequenced reads from all genotypes were trimmed to remove adaptor sequences using cutadapt^56^ v3.4 with the following parameters: -a AGATCGGAAGAGCACACGTCTGAACTCCAGTCA -A AGATCGGAAGAGCGTCGTGTAGGGAAAGAGTGT -q 20 --minimum-length 25. Trimmed reads were aligned to GRCh38 human reference GENCODE release 28 using STAR 1-pass v2.7.8.a^57^ with the following parameters: --outFilterType BySJout --outFilterMultimapNmax 20 --alignSJoverhangMin 8 --alignSJDBoverhangMin 1 --outFilterMismatchNmax 999 -- outFilterMismatchNoverReadLmax 0.04 --alignIntronMin 20 --alignIntronMax 1000000 -- alignMatesGapMax 1000000 --outSAMtype BAM SortedByCoordinate. Quantitation of aligned reads was performed using the featureCounts of the subread package v2.0.1^58^ with annotation sourced from GRCh38 human reference GENCODE release 28 and the following parameters: - s2 -p. After removing outliers, loss-of-function of DLK and LZK genes was analyzed using four vincristine-treated and four untreated biological replicates across the WT, DLK KO, LZK KO, and DLK/LZK double-KO cortical neurons. Loss-of-function of *JUN* and *ATF2* genes was analyzed using 5 vincristine-treated and 5 untreated biological replicates in both control and *JUN* KD cortical neurons, as well as 4 vincristine-treated and 5 untreated ATF2 KD cortical neurons. Differentially expressed genes (DEGs) were analyzed by comparing vincristine-treated cells to untreated cells based on the design ‘∼treatment + 0’ using DESeq2^59^ v1.30.1 in R v4.0.3 (https://cran.r-project.org). Log2-transformed fold changes were computed using the ‘ashr’ shrinkage method^60^ implemented in DESeq2. Genes were considered statistically significant if the false discovery rate (FDR) was below 0.1. The FDR was calculated using Benjamini-Hochberg (BH) method^61^. MA plots were created by plotting the mean normalized read counts on the x-axis and log2 fold changes on the y-axis across genes using ggplot2 in R.

### Lentivirus production

Lentivirus were generated as previously described^52^. In one 10 cm dish, 7 million Lenti-X HEK- 293T cells (Takara Bio, #632180) were seeded in 9 mL DMEM (Gibco, cat. no 11995065) supplemented with 10% FBS (Gibco, #10437028). The next day, a transfection mix was prepared containing 1 µg of Lenti plasmid, 3 µg of third generation packaging mix (1:1:1 mix of three plasmids), 12 µL Lipofectamine 3000 Reagent (ThermoFisher, #L3000008), and 250 µL Opti-MEM I Reduced Serum Medium (GIBCO, #31985070). The mix was vortexed, spun down briefly, incubated at room temperature for 40 minutes, then added dropwise to the HEK cells and gently swirled to mix. The next day, the media was replaced with 18 ml fresh 10% FBS DMEM supplemented with 1:500 ViralBoost (Alstem, #VB100). After two days of incubation, the media was collected into a 50 ml Falcon tube, supplemented with 6 ml Lenti-X Concentrator (Takara Bio; #631231), mixed thoroughly, and stored at 4°C for 48 hours. The supernatant was then spun down at 4°C for 45 minutes at 1,500 x g. The supernatant was aspirated, and the pellet was resuspended in 300 mL of PBS.

### Statistical analysis

All quantification of microscopy images was performed blinded. Fluorescent intensities were quantified unless otherwise stated. Statistical analyses were performed using GraphPad Prism 9. Gene ontology analysis was performed in R version 4.0.3 (2020-10-10). String analysis^62^ was performed with https://string-db.org version 11.5. Venn diagrams were generated using Venny 2.1 (https://bioinfogp.cnb.csic.es/tools/venny).

### Animal protocol

Animal care and experimental procedures were performed in accordance with NICHD protocols 20-003 and 23-003 (Le Pichon lab) approved by the *Eunice Kennedy Shriver* National Institute of Child Health and Human Development ACUC and animal protocol number NEI- 606 (Li lab) approved by the National Eye Institute ACUCC57. C57Bl/6J (Jax#000664) mice of both sexes, aged between 6-10 weeks old, were used in this study.

### Optic nerve crush

Mice were anesthetized with isoflurane, and artificial tears ointment was applied on the non-surgical eye. Crush injury was delivered to the left eye using an intra-orbital approach. The conjunctival layers were incised with a pair of fine scissors and two pairs of #5 fine forceps were used to gently separate the soft tissue in the intra-orbital space behind the eye until the optic nerve was visible. A pair of #5 forceps was used to manually crush the optic nerve for 5 seconds. The eyeball was gently pushed back into the orbit, and analgesia (6 mg/kg Meloxicam) was delivered subcutaneously. Animals were then returned to their home cages and monitored.

### Tissue collection and processing

Animals were injected intraperitoneally and anesthetized with 2.5% Avertin and decapitated. The eyes were removed from the skull and the proximal optic nerve was collected and rapidly frozen on dry ice. The eyes were fixed in 4% paraformaldehyde for 1 hour, and then stored in 1X PBS. Prior to sectioning, the eye was cryopreserved in 30% sucrose. The cornea was removed and the eye cup, including the retina, was frozen in OCT for cryosectioning at 16 µm onto positively charged slides using a Leica CM3050 S Research Cryostat.

### Mild traumatic brain injury (mTBI)

For mTBI experiments, Dlk fl/fl mice were obtained from Dr. Aaron DiAntonio and crossed to Rbp4-Cre mice (Rbp4-Cre (B6.FVB(Cg))-Tg(Rbp4-cre)KL100Gsat/Mmucd, MMRRC Stock No. 037128-UCD) such that the floxed allele would be homozygous. These mice were further crossed to Sun1-sfGFP (B6;129-Gt(ROSA)26Sortm5(CAG-Sun1/sfGFP)Nat/J, Jax Stock No. 021039) such that resulting experimental mice were Rbp4-Cre^+^::Dlk^fl/fl^::Sun1-GFP^+/wt^. Control animals were Rbp4-Cre^+^::Sun1-GFP^+/wt^. Unilateral closed-skull mTBI was administered as previously described^37^ using the Leica Impact One (Leica Biosystems, #39463920) controlled cortical impact (CCI) device. Briefly, Mice receiving injury were anesthetized and the impactor with a 3 mm piston tip mounted on the stereotax at an angle of 10° from the vertical plane was centred roughly at Bregma and moved 2 mm lateral to the midline. The impactor was driven at a velocity of 5 m/s, depth of 1.5 mm, and dwell time of 200 ms. Animals were given 5 mg/kg Meloxicam subcutaneously for analgesia immediately after injury and monitored after removal of anaesthesia to evaluate righting reflex. Mice exhibiting tissue deformation following injury were excluded.

### Immunohistochemistry

Sections were thawed at room temperature for 10 minutes and washed twice with PBS and twice with 0.1% TritonX in PBS (0.1% PBST) for 10 minutes. Sections were blocked for 1 hour at room temperature with 5% normal donkey serum (NDS) in 0.1% PBST. Sections were stained with the selected primary antibody overnight at 4 degrees in 0.5% Donkey serum in 0.1% PBST. Primary antibody was washed four times with 0.1% PBST for 10 minutes at room temperature. Secondary incubation was performed by using the appropriate secondary antibody at a concentration of 1:500 in 0.1% PBST for one hour. Secondary antibody was washed four times with 0.1% PBST for 10 minutes at room temperature. Neurotrace (ThermoFisher, #N21483) labelling was performed by diluting at a concentration of 1:600 in PBS and incubating for 30 minutes. Sections were washed three times with PBS and DAPI was used as a nuclear counterstain. Antibodies used: p-cJun (S63) (CST, #9261S, 1:250), total cJun (CST, #9165, 1:250) and p-ATF2 (T71) (Abcam, #ab32019, 1:250).

### HEK cell culture and transfection

HEK-293A cells were cultured in DMEM (Gibco, #11995065) supplemented with 10% FBS (Gibco; #10437028). For transfection, cells were dissociated using Trypsin-EDTA (0.25%) (ThermoFisher, #25200056) and plated in DMEM, 10%FBS. The day after plating cells were transfected with 1 μg DNA plasmid, 8 μl lipofectamine 2000 (Invitrogen, #11668030) for 24 hours. Cells were fixed with 4% cold PFA for 10 minutes, washed once with PBS and immunocytochemistry was performed.

### ChIP-seq

ChIP-seq was done as previously described^63^. Briefly, Cells were dissociated using Accutase, scraped off the plate, and transferred to a conical tube. After two PBS washes, cells were resuspended in 9.375 ml of BrainPhys media. To fix the cells, 625 µL of 16% formaldehyde was added to achieve a final concentration of 1% and incubated for 10 minutes at 37°C. The reaction was quenched by adding 1.42 M glycine, mixing by inversion, and incubating for 5 minutes at room temperature. Cells were pelleted by centrifugation at 2000 g for 5 minutes at 4°C and washed twice with cold PBS. Cell pellets were then counted, flash frozen in aliquots of 20M cells per tube, and stored at −80°C. For chromatin immunoprecipitation (ChIP), cell pellets were thawed on ice for 10 minutes. Cells were lysed and sonicated using the truChIP Chromatin Shearing Kit (Covaris 520154) according to the manufacturer’s instructions. Chromatin was sonicated in Covaris milliTUBE 1 ml AFA tubes under the following conditions: 30 minutes total, PIP 75, duty factor 15%, CPB 1000, using a Covaris ME220 sonicator. Sonicated samples were centrifuged at 20,000 g for 15 minutes at 4°C, and the supernatant was collected. Chromatin concentration was determined using the Qubit high sensitivity DNA kit. A 20 µg aliquot of chromatin was used per 1 ml ChIP reaction, adjusted with RIPA buffer. For immunoprecipitation, 30 µl of Protein A/G beads conjugated with 2 µg of antibody (pre-incubated with beads resuspended in 500 µl RIPA for 3 hours) was added, and the mixture was incubated overnight at 4°C on a rotating platform. The beads were then washed once with low-salt wash buffer, twice with high-salt wash buffer, twice with lithium chloride wash buffer, and twice with 1X TE buffer. Chromatin was eluted in 200 µl of freshly prepared direct elution buffer supplemented with 2 µl RNase A (10 mg/ml) and incubated on a thermomixer at 65°C and 800 rpm overnight. The following day, beads were removed, and the supernatant was treated with 3 µl Proteinase K (20 mg/ml) at 55°C and 1200 rpm for 2 hours to reverse crosslinks. DNA was purified using the Zymo DNA Clean and Concentrator kit (Zymo, D4033) and eluted in 20 µl deionized water. For library preparation, 10 µl of purified DNA was used per sample. The DNA was prepared with Template Preparation D Buffer (2 µl) and Template Preparation D Enzyme (1 µl) and subjected to end repair and A-tailing in a thermocycler. Library synthesis was performed by adding Library Synthesis D Buffer (1 µl) and Library Synthesis D Enzyme (1 µl) followed by library amplification using Library Amplification D Buffer (25 µl), Library Amplification D Enzyme (1 µl), Nuclease-free water (4 µl), and an Illumina-compatible indexed primer (5 µl). Amplification consisted of an initial extension at 72°C for 3 minutes, denaturation at 98°C for 2 minutes, followed by 4 cycles of 98°C for 20 seconds, 67°C for 20 seconds, and 72°C for 10 seconds for index addition, and 12 cycles of 98°C for 20 seconds and 72°C for 10 seconds for amplification. Libraries were cleaned using a SPRIselect bead double size selection and sequenced on a NovaSeq platform using PE50.

### ChIP-seq analysis

Analysis of ChIPseq data was done as described in^64^. Reads were aligned with Bowtie2 (version 2.5.3)^65^ to the hg38 genome, with the following parameters (-N 1 --local --very-sensitive-local -- no-unal --no-mixed --no-discordant --phred33 -I 10 -X 700 -x). Reads that aligned to the hg38 encode blacklist regions were removed using Samtools 1.19^66^. Duplicate reads were removed with picard (version 3.2.0 broadinstitute.github.io/picard/). Samples that were sequenced in multiple runs were merged before duplicate removal. Peak calling was done using MACS (version 2.2.7.1)^67^ after merging biological replicates and using IgG as a control. For visualization, biological replicate signal was merged into a single bigwig file. These files were created using Deeptools (v3.5.5)^68^ with the bamCoverage function using RPKM normalization, bin size 5. For analyzing differential binding between, we used Diffbind (DiffBind 3.14.0)^69^ . A consensus peakset was created using the peaks identified with the two conditions to be compared and differentially enriched peaks were identified using the DESeq2 method. Signal was analyzed using the summit method and a 400bp bin centered on the summit. Peaks were considered as differentially enriched when the log2FC was larger than one and adjusted p value smaller than 0.05. Heatmaps representing signal strength were generated using deeptools on the peaks that showed no significant difference between JUN and ATF2, peaks with more enrichment of JUN, and peaks with higher enrichment of ATF2. Heatmaps comparing the effect of vincristine and phosphorylation mutants were generated by plotting the signal on a file containing a merge of all peaks identified in either all JUN samples or all ATF2 samples. To assign peaks to genes we used GREAT analysis (v4.0.4)^70^ using the same list of peaks used for the Diffbind. The whole hg38 genome was used as background and Basal plus extension association rule was used with the default 5kb upstream, 1kb downstream setting. Extension distance was changed from 1Mb to 50kb for higher stringency.

## Author Contributions

Conceptualization: J.G.-D. and C.E.L.P.

All experiments using i^3^ Neurons: J.G.-D, M.N., and A.L.S.

ChIPseq: J.G.-D, Z.Z. and P.P.R.

CRISPRi screen: J.G.-D, L.H.E.T.

RNAseq: J.G.-D, M.S., and R.D. mTBI: M.R.A and E.Y.H.L.

Optic nerve crush: J.G.-D., M.R.A, J.W., F.N.-N, P.M.H and N.A.

Writing manuscript: J.G.-D. and C.E.L.P.

Editing manuscript: all authors. Visualization: J.G.-D. and C.E.L.P.

Supervision: R.D., W.L., K.C-C.C., P.P.R., J.G.-D. and C.E.L.P.

Funding acquisition: C.E.L.P., W.L., K.C-C.C., and P.P.R.

## Acknowledgements

We would like to thank Dr. Vincent Schram and the NICHD Microscopy Core, Drs. Tianwei Li and James Iben and the NICHD Molecular Genomics Core, Drs. Martha Kirby and Stacie Anderson and the NHGRI FACS Core. We would also like to thank Dr. Michael Ward and the members of his lab for advice.

## Data Availability

All sequencing data have been deposited at GEO GSE295687 (RNAseq) and GEO: GSE282354 (ChIP seq) and will be made publicly available upon publication. CRISPRi screen data are provided in Supplemental table 2. All other raw data reported in this paper will be shared by the lead contact upon request.

---

Supplemental Figure 1:

A. Schematic representation of DLK (MAP3K12) knockout (KO) strategy. WT i^3^ iPSCs were transfected with two gRNAs targeting *DLK* exons 3-5. Primers used for knockout validation flanking *DLK* exons 3-5 are shown in red.

B. Schematic representation of LZK (MAP3K13) knockout (KO) strategy. WT i^3^ iPSCs were transfected with two gRNAs targeting *DLK* exon 8. Primers used for knockout validation flanking *LZK* exon 8 are shown in red.

C. PCR of WT DLK, LZK and double DLK/LZK KOs using DLK KO primers and LZK KO primers in cortical neuron iPSCs.

D. PCR of WT DLK, LZK and double DLK/LZK KOs using DLK KO primers and LZK KO primers in sensory neuron iPSCs.

E. Representative Western blots of WT, DLK, LZK and double DLK/LZK KO cortical i^3^Neurons. Immunoblot for DLK and loading control b-actin.

F. Representative Western blots of WT, DLK, LZK and double DLK/LZK KO sensory i^3^Neurons. Immunoblot for DLK and loading control b-actin.

G. Representative Western blots of WT, DLK, LZK and double DLK/LZK KO cortical i^3^Neurons untreated (UT) and treated with 5 nM vincristine for 24 hours. Immunoblot for

γH2AX and loading control GAPDH.

H. Quantification of relative γH2AX levels after 5 nM vincristine for 24 hours in WT, DLK, LZK and double DLK/LZK KO cortical i^3^Neurons. Results normalized to untreated (UT). Results are represented as mean ± SEM. N=4 independent differentiations. Two-way ANOVA, Bonferroni correction, p<0.005 ***, p<0.001 ****.

Supplemental Figure 2:

A. Schematic representation of the addition of the transcription inhibitor actinomycin D (ActD) 2, 4, 8, and 24 hours after treatment with vincristine.

B. Quantification of the relative viability of cortical i^3^Neurons after 48 hours of 5 nM vincristine and ActD at different time points after vincristine treatment. N=3 independent differentiations. One-way ANOVA, Bonferroni correction, ns = not significant, p<0.01

**, p<0.005 ***.

C. Number of differentially expressed genes in WT, DLK, LZK and double DLK/LZK KO cortical i^3^Neurons 16 hours after 5nM vincristine treatment.

D. Dot plot showing top 10 enriched biological process gene ontology analysis categories for genes enriched in cortical i^3^Neurons 16 hours after 5nM vincristine treatment.

E. Positive regulation of apoptosis gene ontology analysis -Log10 false discovery rate (FDR) comparison of genes enriched after vincristine treatment in WT, DLK, LZK and double DLK/LZK KO cortical i^3^Neurons.

F. Response to ER stress gene ontology analysis -Log10 false discovery rate (FDR) comparison of genes enriched after vincristine treatment in WT, DLK, LZK and double DLK/LZK KO cortical i^3^Neurons.

G. Venn diagram showing the common genes regulated by DLK and LZK after vincristine treatment.

Supplemental Figure 3:

A. Dot plot showing top 10 enriched biological process gene ontology analysis categories for genes identified to protect neurons from vincristine-induced neurodegeneration in i^3^Neurons.

B. Representative western blots of control, JUN and ATF2 gRNA cortical i^3^Neurons. Immunoblot for ATF2, cJun and loading control b-actin.

C. Quantification of normalized ATF2 levels in control, JUN and ATF2 gRNA cortical i^3^Neurons. Results normalized to Control gRNA. Results represented as mean ± SEM. N=3 independent differentiations. One-way ANOVA, Bonferroni correction, p<0.0001

****.

D. Quantification of normalized cJun levels in Control, JUN and ATF2 gRNA cortical i^3^Neurons. Results normalized to control gRNA. Results are represented as mean ± SEM. N=3 independent differentiations. One-way ANOVA, Bonferroni correction, p<0.05 *.

Supplemental Figure 4:

A. Representative images of WT, DLK, LZK and double DLK/LZK KO cortical i^3^Neurons 24 hours after 5 nM vincristine treatment. Immunostaining for p-cJun S63 (green), DAPI (blue). Scalebar 15 µm.

B. Quantification of relative nuclear p-cJun S63 levels UT and after 5 nM vincristine for 24 hours in WT, DLK, LZK and double DLK/LZK KO cortical i^3^Neurons. Results normalized to WT UT. Results are represented as mean ± SEM. N=3 independent differentiations. Two-way ANOVA, Bonferroni correction, p<0.05 *, p<0.01 **, p<0.005

***, p<0.001 ****.

C. Quantification of relative nuclear total cJun levels UT and after 5 nM vincristine for 24 hours in WT, DLK, LZK and double DLK/LZK KO cortical i^3^Neurons. Results normalized to WT UT. Results represented as mean ± SEM. N=3 independent differentiations. Two-way ANOVA, Bonferroni correction, p<0.01 **, p<0.005 ***.

D. Normalized JUN RNA levels UT and after 5 nM vincristine (vinc) for 16 hours in WT, DLK, LZK and double DLK/LZK KO cortical i^3^Neurons.

E. Representative Western blots of control and JUN gRNA i^3^Neurons transduced with EGFP, WT cJun and cJun S63A S73A (cJun AA)-expressing lentivirus. Immunoblot for cJun and loading control b-actin.

F. Quantification of the relative viability of JUN gRNA cortical i^3^Neurons UT or transduced with EGFP, WT cJun and cJun S63A S73A (cJun AA)-expressing lentivirus after 48 hours of 5 nM vincristine. Results represented as mean ± SEM. N=3 independent differentiations. Two-way ANOVA, Bonferroni correction, ns = not significant, p<0.01

**, p<0.005 ***.

Supplemental Figure 5:

A. Representative images of WT cortical i^3^Neurons UT and 24 hours after 5 nM vincristine treatment. Immunostaining for p-P38 (green), p-cJun S63 (orange) and DAPI (blue). Scalebar 20 µm.

B. Quantification of relative nuclear p-P38 levels UT and after 5 nM vincristine for 24 hours in WT cortical i^3^Neurons. Results normalized to WT UT and represented as mean ± SEM. N=3 independent differentiations. Two-way ANOVA, Bonferroni correction, ns = not significant.

C. Quantification of the relative viability of cortical i^3^Neurons after 48 hours of 5 nM vincristine, 5 nM vincristine + 100 nM P38 inhibitor. N=3 independent differentiations. Two-way ANOVA, Bonferroni correction, ns = not significant.

Supplemental Figure 6:

A. Representative images ipsilateral and contralateral laver V neurons in WT and DLK cKO mice showing layer V GFP+ nuclei. Immunostaining for DAPI (blue), sfGFP (green) and pATF2 (orange).

B. Representative isodensity maps display the topological survival of Brn3a+RGCs in uninjured retinas of control and ATF2 gRNA mice. ATF2 knockdown does not induce degeneration in uninjured retinas. Colorscale for isodensity maps ranges from 0 (purple) to 5800 (red) RGCs/mm^2^.

C. Quantification of percentage Brn3a-positive RGCs in the retinas of control and ATF2 gRNA mice at in uninjured retinas. N = 5-6 mice per condition. Unpaired t-test, ns = not significant.

D. Quantification of nuclear ATF2 levels in Rbpms +ve cells in control and ATF2 gRNA retinas 3 weeks after gRNA transduction.

Supplemental Figure 7:

A. MA plots showing the transcriptional changes in untreated (UT) ATF2 gRNA i^3^Neurons compared to control gRNA. Genes that are significantly differentially expressed are represented by red dots.

B. MA plots showing the transcriptional changes in untreated (UT) JUN gRNA i^3^Neurons compared to control gRNA. Genes that are significantly differentially expressed are represented by red dots.

C. Representative western blots of Control and ATF2 gRNA i^3^Neurons transduced with EGFP, WT ATF2 and ATF2 T69A T71A (ATF2 AA)-expressing lentivirus for 72 hours. Immunoblot for ATF2, cJun and loading control b-actin.

D. Quantification of the relative total cJun levels in control and ATF2 gRNA i^3^Neurons untreated (UT) or transduced with FLAG-ATF2 and FLAG-ATF2 T69A T71A (ATF2 AA)-expressing lentivirus for 72 hours.

E. Heatmap showing ChIP-seq signal at sites identified as significantly stronger in FLAG- ATF2 binding compared to JUN-FLAG (top cluster), no change in binding (middle cluster) and weaker binding (bottom cluster). Signal was centered on the summit of the peaks.

MA plot comparing the differences in binding between FLAG-ATF2 and JUN-FLAG. Peaks with a Log2 FC larger than 1 and an adjusted p lower than 0.05 were identified as significantly differentially enriched and labelled in pink.

