## Supplemental for "ATF2 phosphorylation is a core transcriptional driver of neuron apoptosis"

Supplemental Figure 1

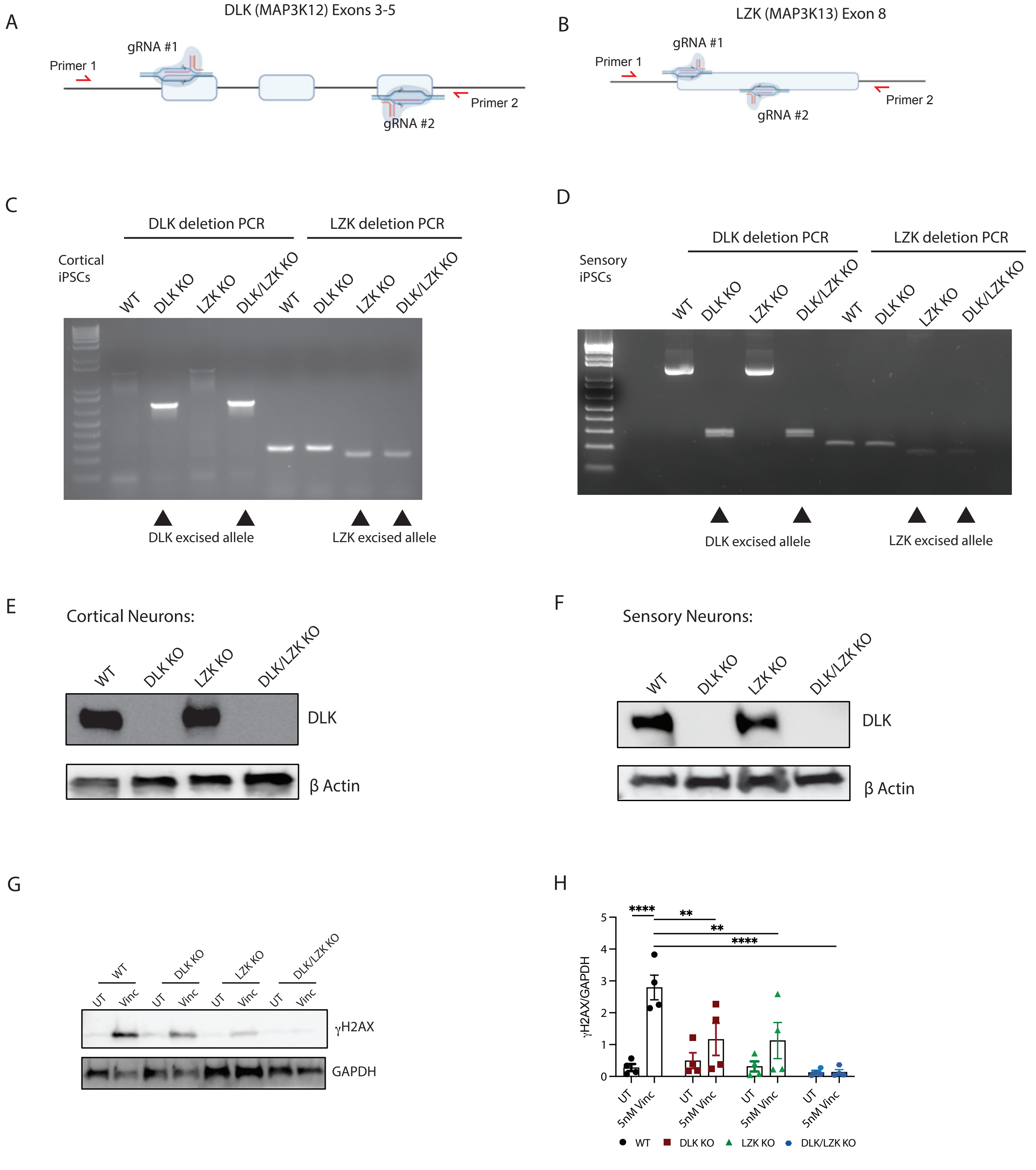

Supplemental Figure 1:

A. Schematic representation of DLK (MAP3K12) knockout (KO) strategy. WT $i^3$ iPSCs were transfected with two gRNAs targeting *DLK* exons 3-5. Primers used for knockout validation flanking *DLK* exons 3-5 are shown in red.
B. Schematic representation of LZK (MAP3K13) knockout (KO) strategy. WT $i^3$ iPSCs were transfected with two gRNAs targeting *DLK* exon 8. Primers used for knockout validation flanking *LZK* exon 8 are shown in red.
C. PCR of WT DLK, LZK and double DLK/LZK KOs using DLK KO primers and LZK KO primers in cortical neuron iPSCs.
D. PCR of WT DLK, LZK and double DLK/LZK KOs using DLK KO primers and LZK KO primers in sensory neuron iPSCs.
E. Representative Western blots of WT, DLK, LZK and double DLK/LZK KO cortical $i^3$Neurons. Immunoblot for DLK and loading control β-actin.
F. Representative Western blots of WT, DLK, LZK and double DLK/LZK KO sensory $i^3$Neurons. Immunoblot for DLK and loading control β-actin.
G. Representative Western blots of WT, DLK, LZK and double DLK/LZK KO cortical $i^3$Neurons untreated (UT) and treated with 5 nM vincristine for 24 hours. Immunoblot for γH2AX and loading control GAPDH.
H. Quantification of relative γH2AX levels after 5 nM vincristine for 24 hours in WT, DLK, LZK and double DLK/LZK KO cortical $i^3$Neurons. Results normalized to untreated (UT). Results are represented as mean ± SEM. N=4 independent differentiations. Two-way ANOVA, Bonferroni correction, $p<0.005$ ***, $p<0.001$ ****.

Supplemental Figure 2

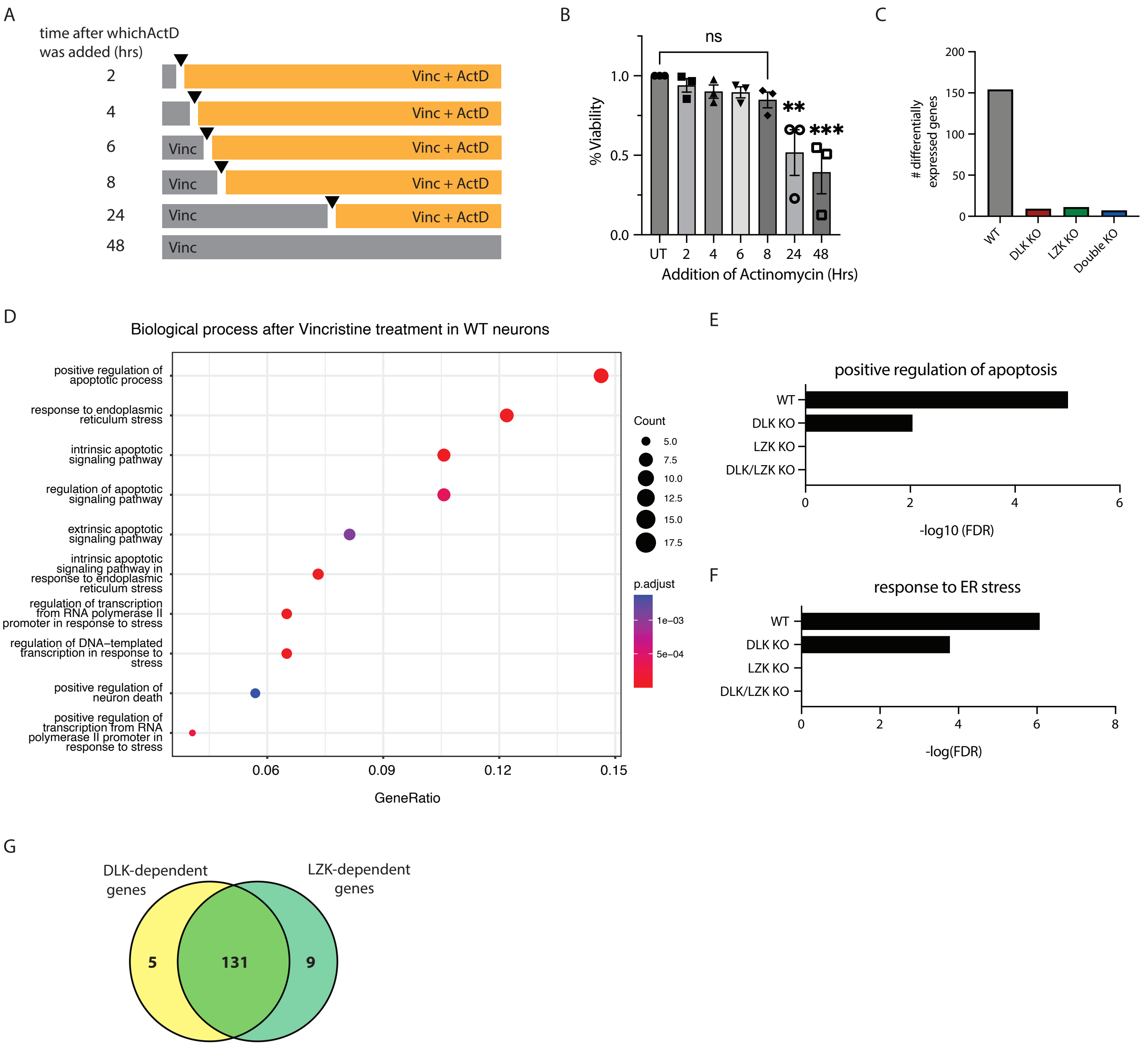

Supplemental Figure 2:

A. Schematic representation of the addition of the transcription inhibitor actinomycin D (ActD) 2, 4, 8, and 24 hours after treatment with vincristine.
B. Quantification of the relative viability of cortical $i^3$Neurons after 48 hours of 5 nM vincristine and ActD at different time points after vincristine treatment. N=3 independent differentiations. One-way ANOVA, Bonferroni correction, ns = not significant, p<0.01 **, p<0.005 ***.
C. Number of differentially expressed genes in WT, DLK, LZK and double DLK/LZK KO cortical $i^3$Neurons 16 hours after 5nM vincristine treatment.
D. Dot plot showing top 10 enriched biological process gene ontology analysis categories for genes enriched in cortical $i^3$Neurons 16 hours after 5nM vincristine treatment.
E. Positive regulation of apoptosis gene ontology analysis -Log10 false discovery rate (FDR) comparison of genes enriched after vincristine treatment in WT, DLK, LZK and double DLK/LZK KO cortical $i^3$Neurons.
F. Response to ER stress gene ontology analysis -Log10 false discovery rate (FDR) comparison of genes enriched after vincristine treatment in WT, DLK, LZK and double DLK/LZK KO cortical $i^3$Neurons.
G. Venn diagram showing the common genes regulated by DLK and LZK after vincristine treatment.

Supplemental Figure 3

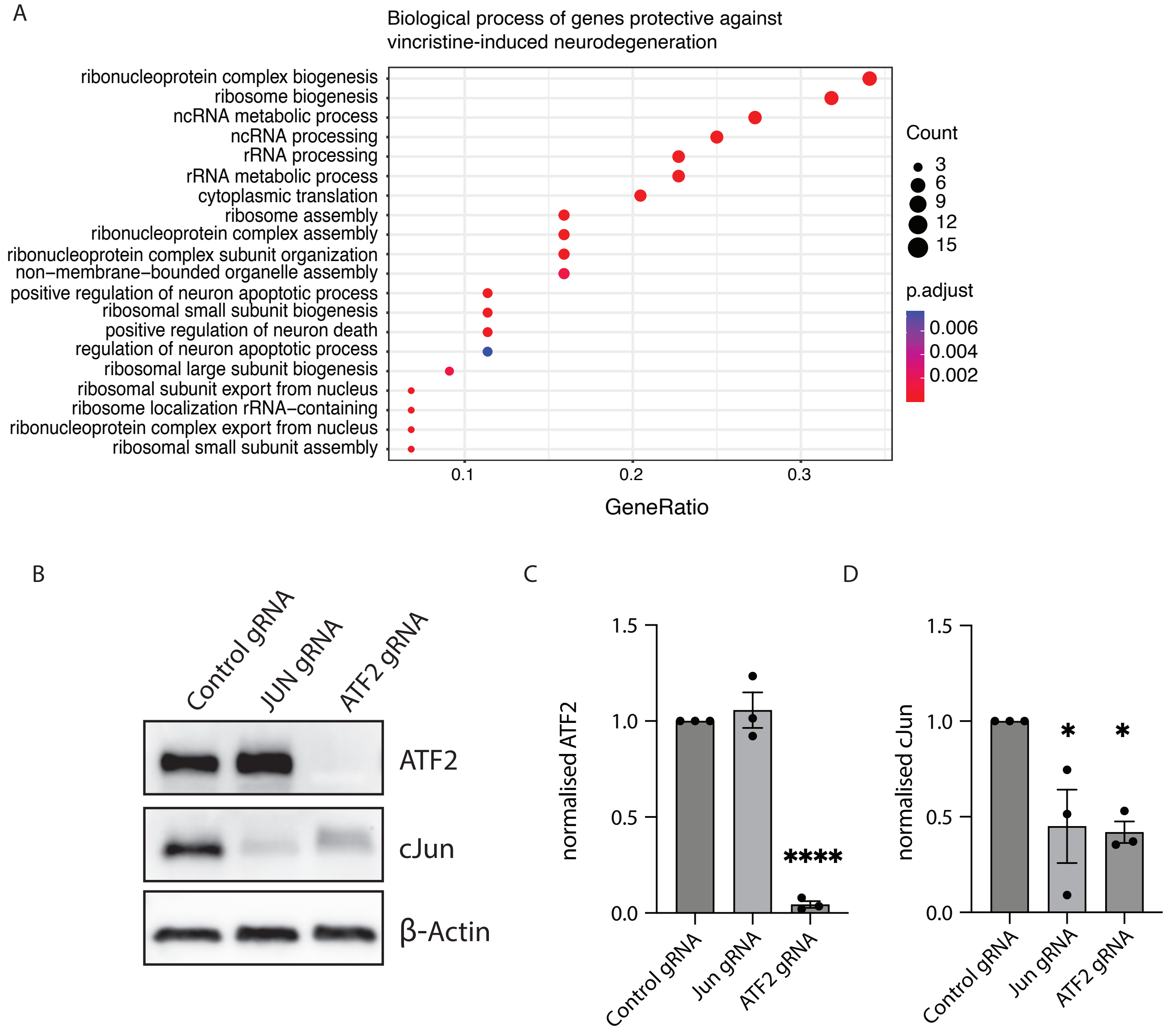

Supplemental Figure 3:

A. Dot plot showing top 10 enriched biological process gene ontology analysis categories for genes identified to protect neurons from vincristine-induced neurodegeneration in $i^3$Neurons.
B. Representative western blots of control, JUN and ATF2 gRNA cortical $i^3$Neurons. Immunoblot for ATF2, cJun and loading control $\beta$-actin.
C. Quantification of normalized ATF2 levels in control, JUN and ATF2 gRNA cortical $i^3$Neurons. Results normalized to Control gRNA. Results represented as mean ± SEM. N=3 independent differentiations. One-way ANOVA, Bonferroni correction, $p<0.0001$ ****.
D. Quantification of normalized cJun levels in Control, JUN and ATF2 gRNA cortical $i^3$Neurons. Results normalized to control gRNA. Results are represented as mean ± SEM. N=3 independent differentiations. One-way ANOVA, Bonferroni correction, $p<0.05$ *.

Supplemental Figure 4

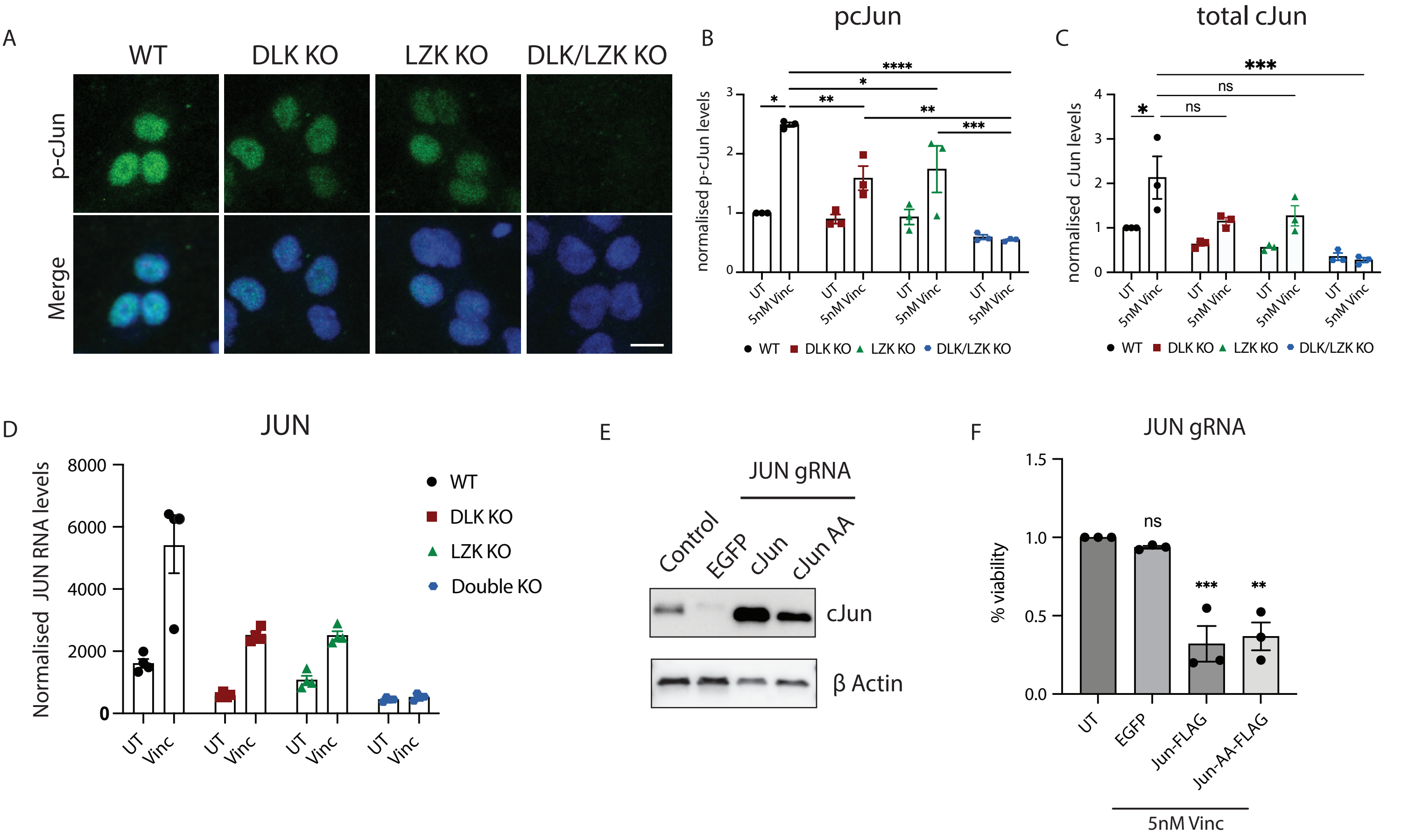

Supplemental Figure 4:

A. Representative images of WT, DLK, LZK and double DLK/LZK KO cortical $i^3$Neurons 24 hours after 5 nM vincristine treatment. Immunostaining for p-cJun S63 (green), DAPI (blue). Scalebar 15 µm.
B. Quantification of relative nuclear p-cJun S63 levels UT and after 5 nM vincristine for 24 hours in WT, DLK, LZK and double DLK/LZK KO cortical $i^3$Neurons. Results normalized to WT UT. Results are represented as mean ± SEM. N=3 independent differentiations. Two-way ANOVA, Bonferroni correction, p<0.05 *, p<0.01 **, p<0.005 ***, p<0.001 ****.
C. Quantification of relative nuclear total cJun levels UT and after 5 nM vincristine for 24 hours in WT, DLK, LZK and double DLK/LZK KO cortical $i^3$Neurons. Results normalized to WT UT. Results represented as mean ± SEM. N=3 independent differentiations. Two-way ANOVA, Bonferroni correction, p<0.01 **, p<0.005 ***.
D. Normalized JUN RNA levels UT and after 5 nM vincristine (vinc) for 16 hours in WT, DLK, LZK and double DLK/LZK KO cortical $i^3$Neurons.
E. Representative Western blots of control and JUN gRNA $i^3$Neurons transduced with EGFP, WT cJun and cJun S63A S73A (cJun AA)-expressing lentivirus. Immunoblot for cJun and loading control $\beta$-actin.
F. Quantification of the relative viability of JUN gRNA cortical $i^3$Neurons UT or transduced with EGFP, WT cJun and cJun S63A S73A (cJun AA)-expressing lentivirus after 48 hours of 5 nM vincristine. Results represented as mean ± SEM. N=3 independent differentiations. Two-way ANOVA, Bonferroni correction, ns = not significant, p<0.01 **, p<0.005 ***.

Supplemental Figure 5

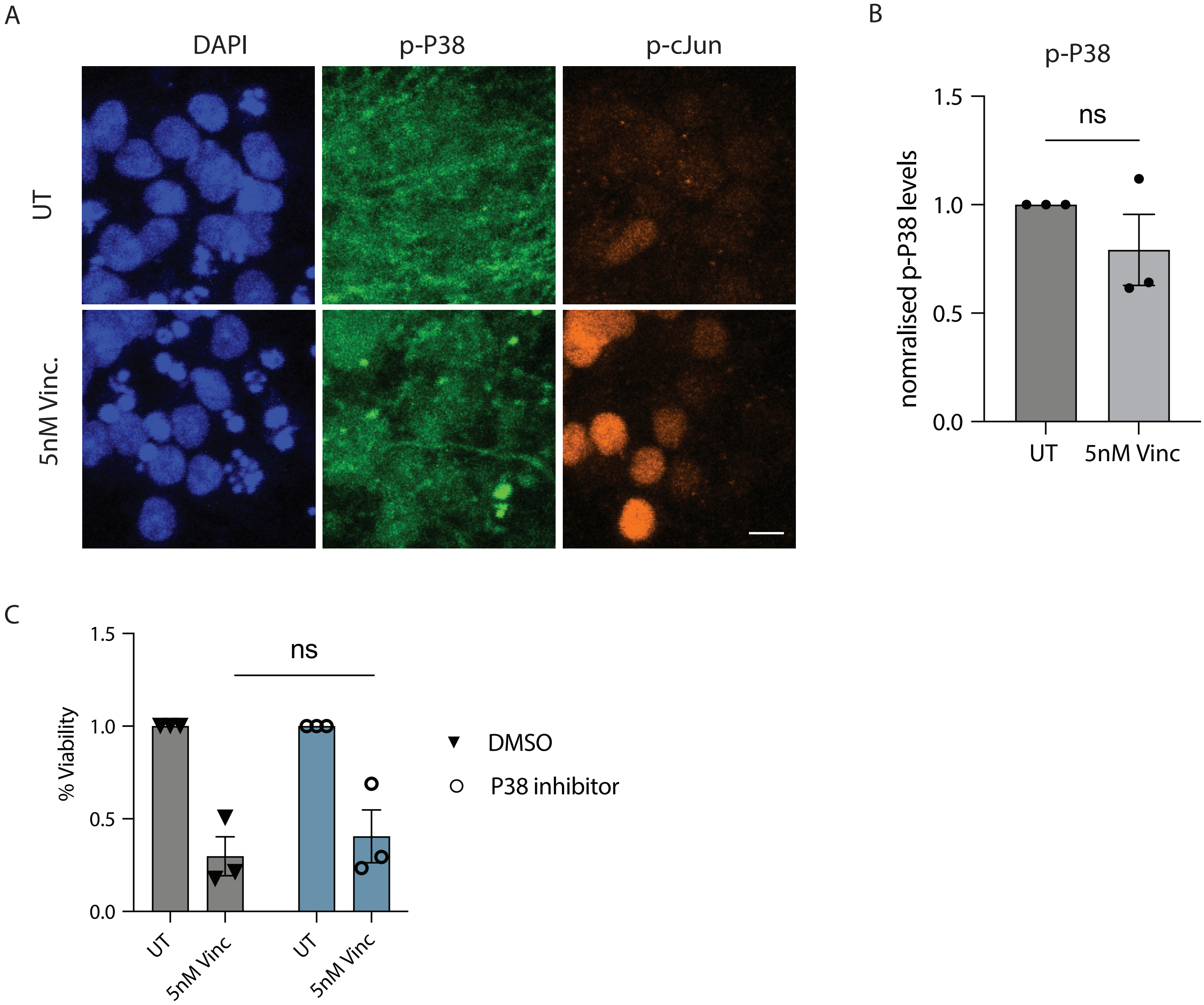

Supplemental Figure 5:

A. Representative images of WT cortical i$^3$Neurons UT and 24 hours after 5 nM vincristine treatment. Immunostaining for p-P38 (green), p-cJun S63 (orange) and DAPI (blue). Scalebar 20 µm.
B. Quantification of relative nuclear p-P38 levels UT and after 5 nM vincristine for 24 hours in WT cortical i$^3$Neurons. Results normalized to WT UT and represented as mean ± SEM. N=3 independent differentiations. Two-way ANOVA, Bonferroni correction, ns = not significant.
C. Quantification of the relative viability of cortical i$^3$Neurons after 48 hours of 5 nM vincristine, 5 nM vincristine + 100 nM P38 inhibitor. N=3 independent differentiations. Two-way ANOVA, Bonferroni correction, ns = not significant.

Supplemental Figure 6

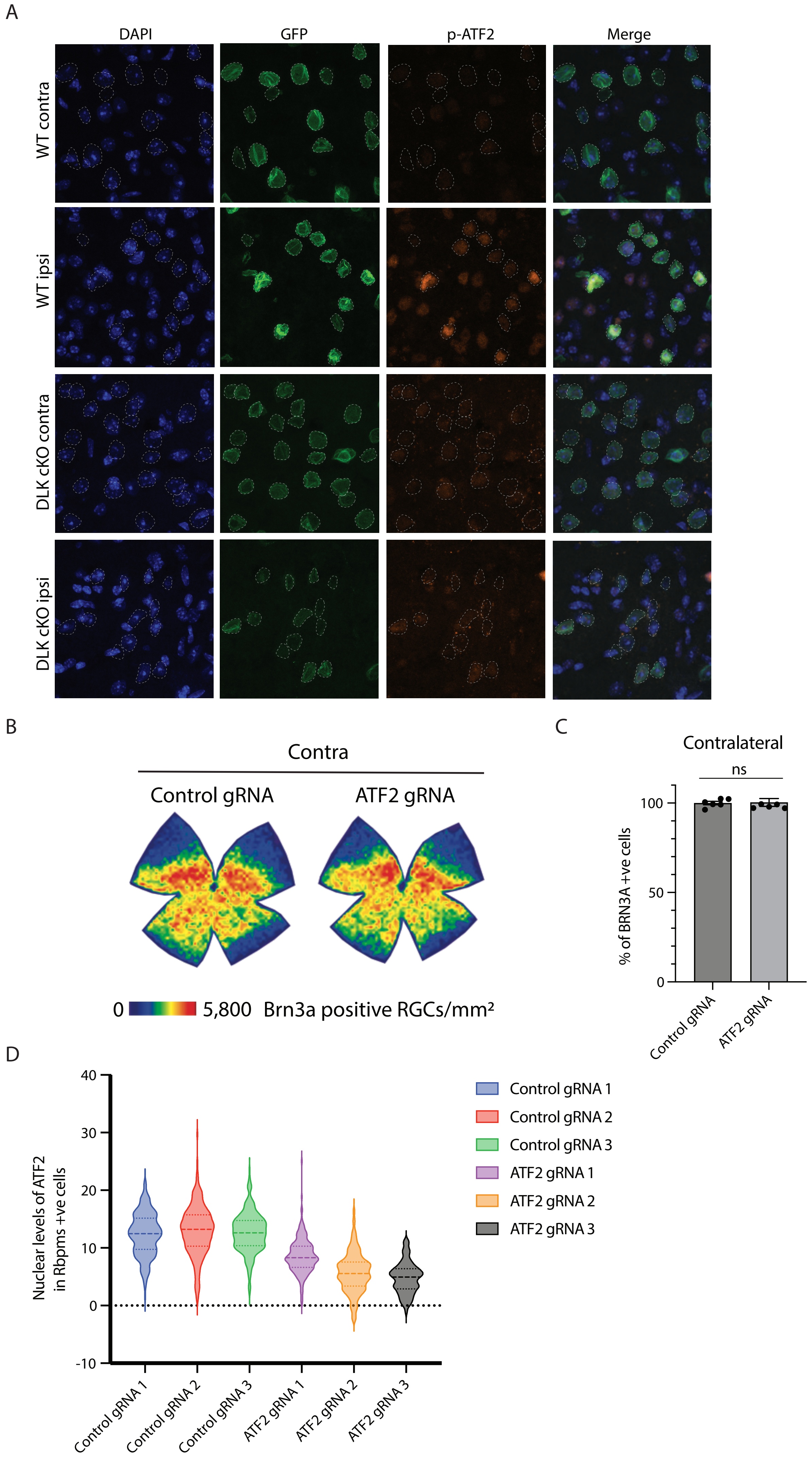

Supplemental Figure 6:

A. Representative images ipsilateral and contralateral laver V neurons in WT and DLK cKO mice showing layer V GFP+ nuclei. Immunostaining for DAPI (blue), sfGFP (green) and pATF2 (orange).
B. Representative isodensity maps display the topological survival of Brn3a+RGCs in uninjured retinas of control and ATF2 gRNA mice. ATF2 knockdown does not induce degeneration in uninjured retinas. Colorscale for isodensity maps ranges from 0 (purple) to 5800 (red) RGCs/mm$^2$.
C. Quantification of percentage Brn3a-positive RGCs in the retinas of control and ATF2 gRNA mice at in uninjured retinas. N = 5-6 mice per condition. Unpaired t-test, ns = not significant.
D. Quantification of nuclear ATF2 levels in Rbpms +ve cells in control and ATF2 gRNA retinas 3 weeks after gRNA transduction.

Supplemental Figure 7

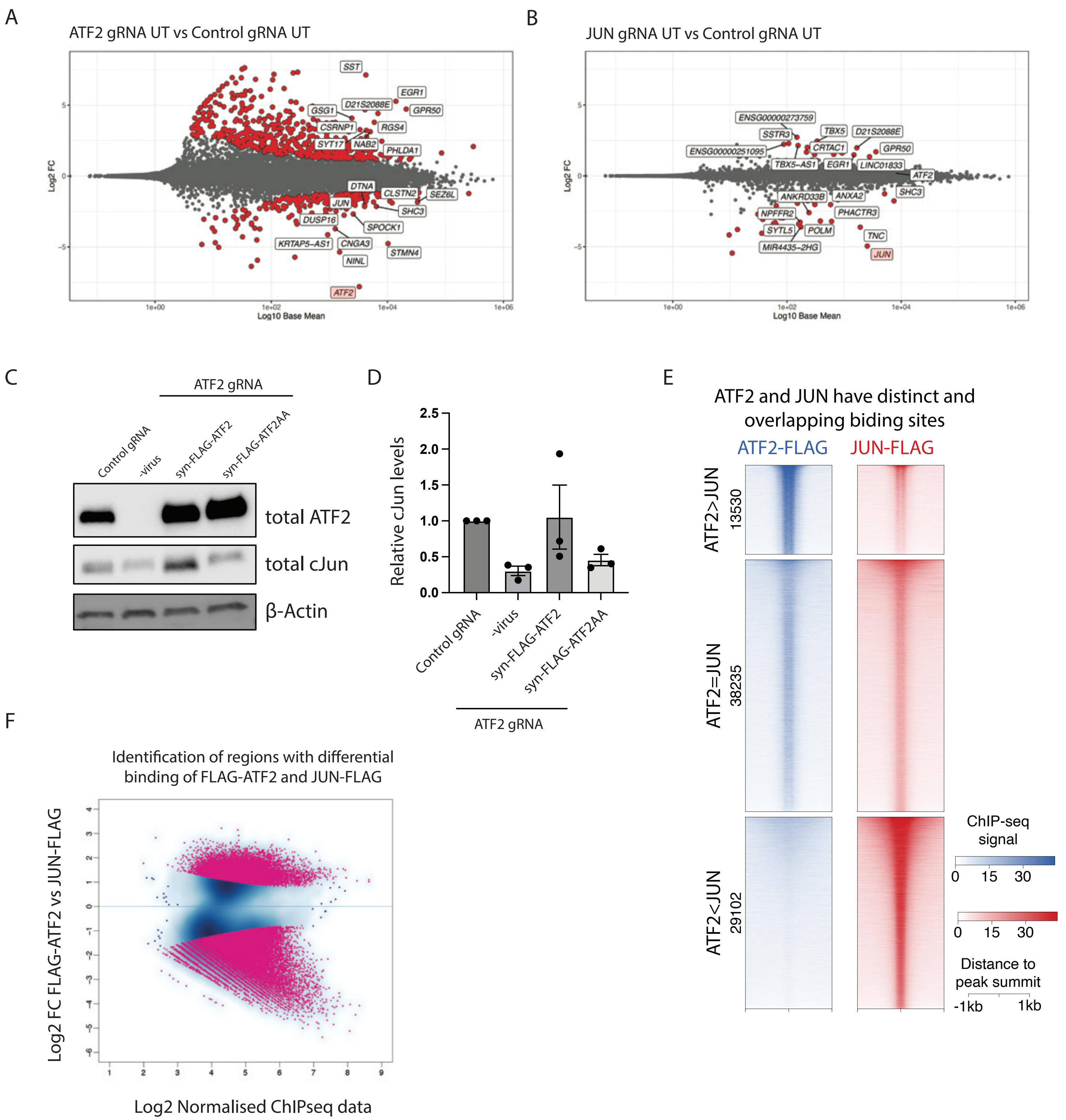

Supplemental Figure 7:

A. MA plots showing the transcriptional changes in untreated (UT) ATF2 gRNA $i^3$Neurons compared to control gRNA. Genes that are significantly differentially expressed are represented by red dots.
B. MA plots showing the transcriptional changes in untreated (UT) JUN gRNA $i^3$Neurons compared to control gRNA. Genes that are significantly differentially expressed are represented by red dots.
C. Representative western blots of Control and ATF2 gRNA $i^3$Neurons transduced with EGFP, WT ATF2 and ATF2 T69A T71A (ATF2 AA)-expressing lentivirus for 72 hours. Immunoblot for ATF2, cJun and loading control $\beta$-actin.
D. Quantification of the relative total cJun levels in control and ATF2 gRNA $i^3$Neurons untreated (UT) or transduced with FLAG-ATF2 and FLAG-ATF2 T69A T71A (ATF2 AA)-expressing lentivirus for 72 hours.
E. Heatmap showing ChIP-seq signal at sites identified as significantly stronger in FLAG-ATF2 binding compared to JUN-FLAG (top cluster), no change in binding (middle cluster) and weaker binding (bottom cluster). Signal was centered on the summit of the peaks.
F. MA plot comparing the differences in binding between FLAG-ATF2 and JUN-FLAG. Peaks with a Log2 FC larger than 1 and an adjusted p lower than 0.05 were identified as significantly differentially enriched and labelled in pink.
